# *C. elegans* SMOC-1 interacts with both BMP and glypican to regulate BMP signaling

**DOI:** 10.1101/2023.01.06.523017

**Authors:** Melisa S. DeGroot, Byron Williams, Timothy Y Chang, Maria L. Maas Gamboa, Isabel Larus, J. Christopher Fromme, Jun Liu

## Abstract

Secreted modular calcium binding (SMOC) proteins are conserved matricellular proteins found in organisms from *C. elegans* to humans. SMOC homologs characteristically contain one or two extracellular calcium (EC) binding domain(s) and one or two thyroglobulin type-1 (TY) domain(s). SMOC proteins in *Drosophila* and Xenopus have been found to interact with cell surface heparan sulfate protein glycans (HSPGs) to exert both positive and negative influences on the conserved bone morphogenetic protein (BMP) signaling pathway. In this study, we used a combination of biochemical, structural modeling, and molecular genetic approaches to dissect the functions of the sole SMOC protein in *C. elegans*. We showed that SMOC-1 binds LON-2/glypican, as well as the mature domain of DBL-1/BMP. Moreover, SMOC-1 can simultaneously bind LON-2/glypican and DBL-1/BMP. The interaction between SMOC-1 and LON-2/glypican is mediated by the EC domain of SMOC-1, while the interaction between SMOC-1 and DBL-1/BMP involves full-length SMOC-1. We further showed that while SMOC-1(EC) is sufficient to promote BMP signaling when overexpressed, both the EC and TY domains are required for SMOC-1 function at the endogenous locus. Finally, when overexpressed, SMOC-1 can promote BMP signaling in the absence of LON-2/glypican. Taken together, our findings led to a model where SMOC-1 functions both negatively in a LON-2-dependent manner and positively in a LON-2-independent manner to regulate BMP signaling. Our work provides a mechanistic basis for how the evolutionarily conserved SMOC proteins regulate BMP signaling.

## INTRODUCTION

The highly conserved bone morphogenetic protein (BMP) pathway is used in a variety of developmental and homeostatic processes across metazoans [1]. BMP signaling is activated upon binding of the secreted BMP ligand to complexes of the type I and type II receptor kinases to induce an intracellular phosphorylation cascade. The signal is transduced by the type II receptor phosphorylating the type I receptor, which in turn phosphorylates the receptor-activated R-Smads. Once phosphorylated, the activated R-Smads complex with common mediator Smads (Co-Smads) and enter the nucleus to regulate downstream gene transcription. Activation of the BMP pathway must be tightly regulated in space, time, level and duration, as mis-regulation of the pathway can cause a variety of disorders in humans, including cancer [2, 3]. Multiple levels of regulatory mechanisms, including at the extracellular level, have been identified to ensure precise control of BMP signaling [4-6]. One class of extracellular regulators of BMP signaling are the secreted modular calcium binding SMOC proteins.

Secreted modular calcium binding proteins (SMOCs) are matricellular proteins belonging to the BM-40/osteonectin/SPARC (basement membrane of 40kDa/secreted protein acidic and rich in cysteine) family [7]. The SPARC family is characterized as containing an extracellular calcium (EC) binding domain and a follistatin-like (FS) domain, and includes related proteins such as BM-40, SMOCs, QR1, SC1/hevin, tsc36/Flik, and testicans [7]. SMOCs are found in metazoans ranging from flies and nematodes to mice and humans. SMOC homologs characteristically contain a thyroglobulin type-1 (TY) domain followed by an EC domain, and in some cases contain a FS domain. Notably, the number and arrangement of the domains vary across species. For example, humans have two SMOC proteins, and both have a FS domain followed by two TY domains and one EC domain [8, 9]. The single *Drosophila* SMOC homolog, Pentagone (Pent), has a FS domain followed by two alternating TY and EC domains [10, 11], while the lone SMOC-1 protein in *C. elegans* has one TY domain followed by one EC domain [12].

Despite the differences in domain structure arrangements, all SMOC proteins that have been studied regulate BMP signaling [10-16]. The underlying mechanistic bases, however, are not identical across organisms. One of the more unifying models from studies of *Drosophila* Pent and Xenopus SMOC-1 suggests that SMOC proteins may function by competing with BMPs for binding to heparan sulfate proteoglycans (HSPGs), thus expanding the range of BMP signaling [17]. However, recent work from *Drosophila* suggests that Pent may have additional unappreciated functions in addition to simply expanding the range of Dpp/BMP (Zhu et al., Dev Cell, 2020). For example, Dally/glypican typically functions in a positive fashion as a BMP co-receptor for BMP signaling in *Drosophila* [18-20]. Yet knocking down enzymes required for the production of HSPGs results in a similar wing disc phenotype as Pent overexpression in the *Drosophila* wing imaginal discs [21], suggesting that the regulatory relationship between these factors is complex. Similarly, Xenopus SMOC-1 is known to antagonize BMP signaling by acting downstream of the BMP receptors [16]. Thus additional studies are needed to further elucidate how SMOC proteins may function to both positively and negatively regulate BMP signaling.

*C. elegans* has a well-conserved BMP-like pathway. The core components of the *C. elegans* BMP pathway include the ligand DBL-1, the type I and type II receptors SMA-6 and DAF-4, the R-Smads SMA-2 and SMA-3, and the Co-Smad SMA-4 [22, 23]. This pathway is known to regulate multiple biological processes including body size and mesoderm development [22, 23]. Changes in BMP pathway activation affect body size in a dose dependent manner with reductions in signaling causing a small (Sma) body size, while increased BMP signaling causing a long (Lon) phenotype [22]. BMP signaling also regulates postembryonic mesoderm development. In particular, mutations in the zinc finger transcription factor SMA-9 cause a loss of the two posterior coelomocytes (CCs) due to a fate transformation in the post-embryonic mesoderm or the M-lineage [24]. Mutations in components of the BMP pathway can suppress the M-lineage defects of *sma-9(0)* mutants (Susm), such that resulting worms have proper specification of the two post-embryonic CCs [24, 25]. Both the body size and the Susm phenotypes can be used to assess the functionality of BMP pathway components. In particular, the Susm assay is highly specific and sensitive to altered BMP signaling activity [24, 25].

*C. elegans* also has a single, conserved SMOC protein, SMOC-1, which is known to regulate BMP signaling. We have previously demonstrated that *C. elegans* SMOC-1 promotes BMP signaling in a cell non-autonomous fashion by acting through the BMP ligand in a positive feedback loop [12]. In this study, we implemented an unbiased approach to identify SMOC-1 interacting proteins. We found that, like SMOC proteins in other organisms [10, 17, 26, 27], *C. elegans* SMOC-1 binds to the HSPG LON-2/glypican via its EC domain. Surprisingly, we also found that full-length SMOC-1 can bind to DBL-1/BMP, and that SMOC-1 can mediate the formation of a LON-2-SMOC-1-DBL-1 tripartite complex in vitro. We showed that while SMOC-1(EC) is capable of overstimulating BMP signaling when overexpressed, both SMOC-1(TY) and SMOC-1(EC) are required for full function of SMOC-1 when expressed at endogenous levels. We further showed that overexpression of SMOC-1 is sufficient to further promote BMP signaling in the absence of LON-2/glypican. Collectively, our data support a model where a SMOC-1-dependent glypican-SMOC-BMP complex inhibits BMP signaling, while the SMOC-1-BMP complex promotes BMP signaling.

## RESULTS

### Generate a *C. elegans* strain with SMOC-1 functionally tagged and detectable

To determine how SMOC-1 functions at the molecular level to regulate BMP signaling, we aimed to tag SMOC-1 at the endogenous locus without disrupting its function. We found that endogenously tagging SMOC-1 with GFP at either the N-terminus after the signal peptide or the C-terminus disrupted SMOC-1 function (Figure S1). Instead, adding two copies of the small FLAG tag to the C-terminus of SMOC-1 at the endogenous locus did not affect SMOC-1 function, based on both the body size and the Susm assays of *smoc-1(jj276[smoc-1::2xflag])* worms (Figure 1). The endogenously tagged SMOC-1::2XFLAG protein, however, was not detectable via western blot (Figure 1B). To overcome this problem, we generated two integrated transgenic lines carrying multicopy transgenic arrays that overexpress SMOC-1::2XFLAG, *jjIs5798[smoc-1::2xflag(OE)]* and *jjIs5799[smoc-1::2xflag(OE)]*. Both *jjIs5798* and *jjIs5799* worms exhibit a long phenotype, similar to *jjIs5119[smoc-1(OE)]* worms overexpressing an untagged *smoc-1*, (Figure 1C, [12]). These data, combined with the data on *smoc-1(jj276)* worms, suggest that *jjIs5798* and *jjIs5799* worms contain a fully functional *smoc-1::2xflag* that is simply overexpressed. The overexpressed SMOC-1::2xFLAG protein from *jjIs5798* and *jjIs5799* worms is detectable via western blot (Figure 1B). The ability to express and detect a functionally-tagged SMOC-1 protein *in vivo* allowed us to identify SMOC-1-interacting partners from worm extracts, and to carry out structure-function studies of the SMOC-1 protein in live worms.

**Figure 1.**
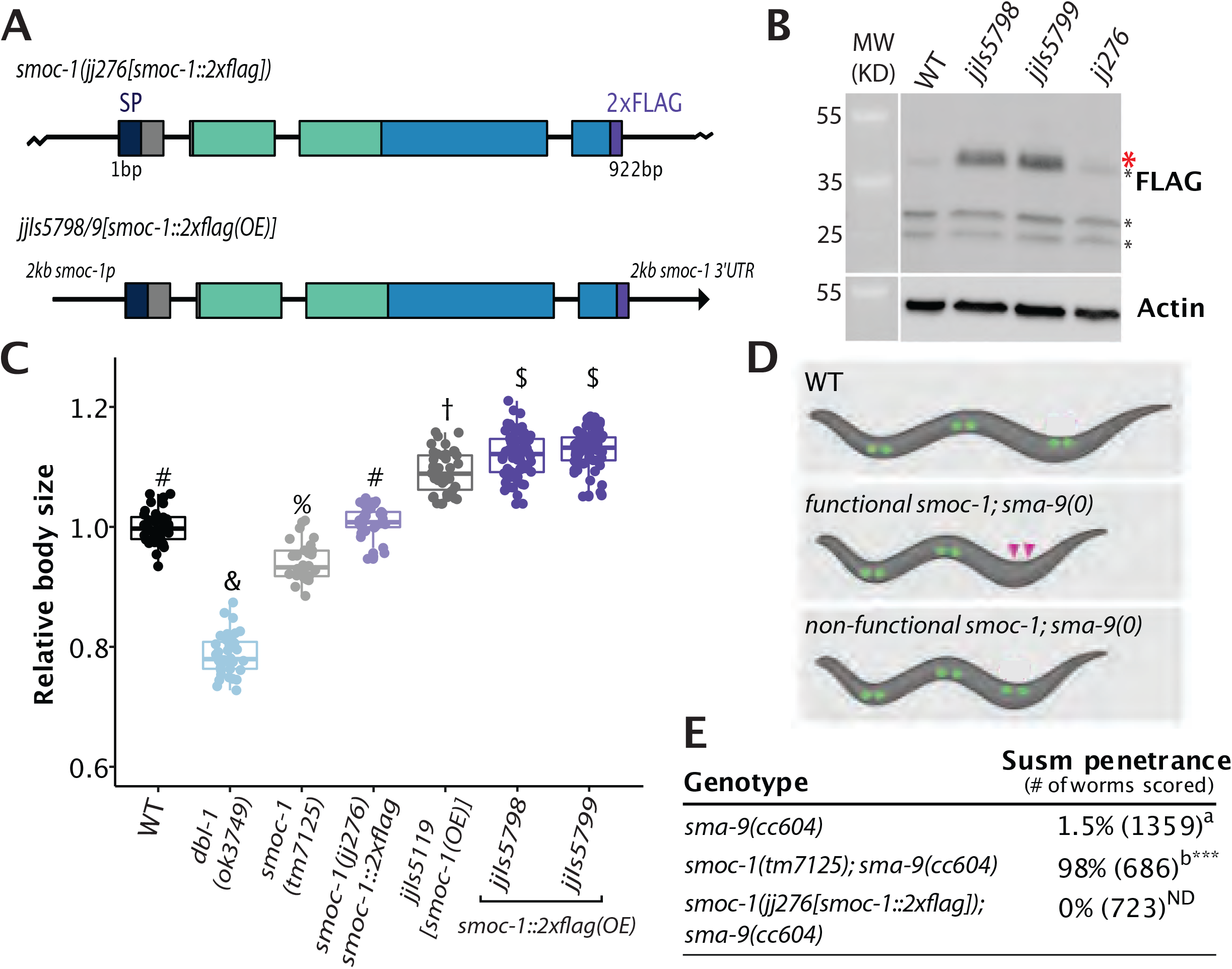
SMOC-1::2xFLAG is fully functional. **A)** Diagrams depicting the *smoc-1::2xflag* endogenous locus (*jj276)* and a transgene with 2kb *smoc-1* promoter and 2kb *smoc-1* 3’UTR flanking the *smoc-1::2xflag* genomic sequence. *jjIs5798* and *jjIs5799* are two integrated multicopy array lines that overexpress *smoc-1::2xflag*. In this and all subsequent figures, protein domains in SMOC-1 are indicated by color: navy, SP, signal peptide; green, TY, thyroglobulin-like domain; blue, EC, extracellular calcium binding domain; purple, 2xFLAG. Exons are represented by colored bars, while introns and intergenic regions are represented by thin black lines. **B)** Western blot of 50 gravid adults of each indicated genotype, probed with anti-FLAG (top) and anti-actin antibodies (bottom). Full length SMOC-1::2xFLAG at ∼41KDa (marked by a red *) is detectable in strains that overexpress SMOC-1::2xFLAG (*jjIs5798* and *jjIs5799*), but not in wild-type (WT) controls nor in *jj276* worms that expresses endogenous SMOC-1::2xFLAG. Black *s indicate non-specific bands detected by the anti-FLAG antibody. **C)** Relative body sizes of various strains at the same developmental stage (WT set to 1.0). *jjIs5119* is an integrated multicopy array line that overexpresses untagged *smoc-1*. Groups marked with distinct symbols are significantly different from each other (*P*<0.001, in all cases when there is a significant difference), while groups with the same symbol are not. Tested using an ANOVA with a Tukey HSD. WT: N=65. *ok3749*: N=38. *tm7125:* N=36. *jj276*: N=31. *jjIs5119*: N=42. *jjIs5798*: N=72. *jjIs5799*: N=68. **D)** Diagrams depicting the Susm phenotype used to test *smoc-1* functionality. Coelomocytes (CC) are represented with green circles, with the two posterior M-derived CCs marked with purple arrowheads. **E)** Table showing the penetrance of the Susm phenotype of various mutant strains. The Susm penetrance refers to the percentage of animals with one or two M-derived CCs as scored using the *arIs37*(*secreted CC::GFP)* reporter (see materials and methods). For each genotype, the Susm data from two independent isolates (see table S3) were combined and presented in the table. ^a^ The lack of M-derived CCs phenotype is not fully penetrant in *sma-9(cc604)* mutants. ^b^ Data from DeGroot et al. [12]. Statistical analysis was conducted by comparing double mutant lines with the *sma-9(cc604)* single mutants. \*\*\**P*<0.001; ND: no difference (unpaired two-tailed Student’s *t*-test).

### SMOC-1 associates with LON-2/glypican in worm lysates and when produced in a heterologous system

We next conducted immunoprecipitation (IP) of whole worm lysate from strains overexpressing SMOC-1::2xFLAG and identified the proteins co-precipitated using mass spectrometry (MS). As controls, we used a strain overexpressing full length SMOC-1 lacking the FLAG tag (hereafter referred to as “untagged SMOC-1”). We conducted two rounds of IP-MS using independent biological samples (see materials and methods). Analysis of the MS results showed that SMOC-1 was detected exclusively in the tagged samples, indicating that the IP via anti-FLAG antibody was highly specific to the FLAG tagged SMOC-1. In both experiments, LON-2/glypican was identified as a strong candidate SMOC-1-interaction partner, as demonstrated by the recovery of 13 and 18 respective peptides that map specifically to LON-2 (Figure 2A, Table S1). No peptides mapping to LON-2 were detected in the untagged SMOC-1 samples.

**Figure 2.**
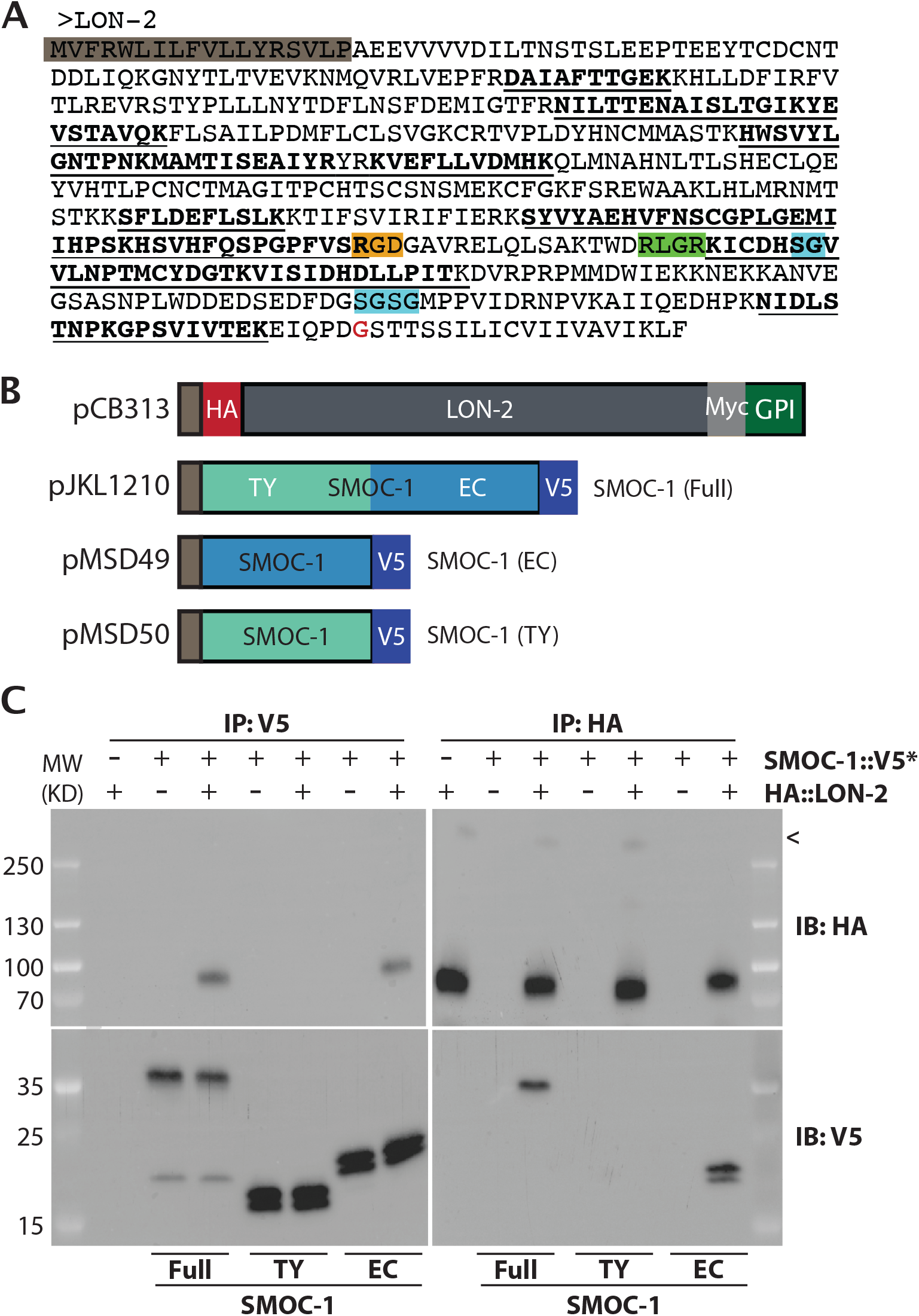
SMOC-1 interacts with LON-2 in worm extracts and *in vitro*. **A)** Sequence of *C. elegans* LON-2 protein with peptide regions detected in the different IP/MS experiments marked. Peptides detected in the full length SMOC-1::2xFLAG pulldown are underlined, while peptides detected in the SMOC-1(EC)::2xFLAG pulldown are in bold. Important regions of LON-2 are also highlighted, including the signal peptide (grey), RGD motif (orange), RLGR consensus putative furin protease recognition site (green), HS GAG attachment sites (aqua), and GPI linkage site (red). **B)** Diagrams of the expression constructs used in the *Drosophila* S2 cell expression system. **C)** Results of co-IP experiments testing the interaction of HA::LON-2 with different versions of SMOC-1::V5* produced in *Drosophila* S2 cells, including SMOC-1(Full)::V5, SMOC-1(TY)::V5, and SMOC-1(EC)::V5. Immunoprecipitation (IP) with anti-V5 beads or anti-HA beads, immunoblot (IB) with anti-HA or anti-V5 antibodies, as indicated. Experiments were independently repeated in triplicate, with representative results shown in this figure. < points to faint bands that may represent glycosylated LON-2. We do not know whether posttranslational modification or cleavage causes SMOC-1 proteins to run as two bands when expressed in *Drosophila* S2 cells.

To test the interaction between SMOC-1 and LON-2 using an independent assay, we conducted coimmunoprecipitation (co-IP) experiments using *Drosophila* S2 cells that overexpress HA::LON-2 and SMOC-1::V5 (Figure 2B). As shown in Figure 2C, we detected association between SMOC-1 and LON-2 in bidirectional IP experiments: IP of SMOC-1::V5 pulled down HA::LON-2, while IP of HA::LON-2 pulled down SMOC-1::V5.

Taken together, results from our co-IP experiments using both worm extracts and the *Drosophila* S2 cell expression system demonstrated that SMOC-1 interacts with the glypican LON-2.

### SMOC-1 interacts with LON-2/glypican through its EC domain

To identify the specific domain via which SMOC-1 interacts with LON-2, we expressed tagged versions of the SMOC-1 EC domain or the SMOC-1 TY domain in *Drosophila* S2 cells, and tested their interaction with LON-2. Bi-directional co-IP experiments showed that SMOC-1(EC), but not SMOC-1(TY), can interact with LON-2 (Figure 2C). To corroborate these co-IP results, we generated transgenic lines overexpressing SMOC-1(EC)::2xFLAG, and conducted IP-MS experiments using the same condition used for IP-MS experiments with full length SMOC-1::2xFLAG (see materials and methods). Results from these experiments supported an interaction between SMOC-1(EC) and LON-2, as 15 peptides that map specifically to LON-2 were recovered (Figure 2A, Table S1). The co-IP experiments using both worm extracts and the *Drosophila* S2 cells demonstrated that the EC domain of SMOC-1 interacts with LON-2/glypican.

### The EC domain of SMOC-1 is sufficient to promote BMP signaling when overexpressed

We then tested the functionality of the SMOC-1 TY and EC domain individually by generating transgenic lines overexpressing either the TY domain or the EC domain tagged with 2xFLAG. We found that when overexpressed, SMOC-1(EC) rescued the body size defect of *smoc-1(0)* animals, making the worms longer than wild-type (WT) worms, although not as long as worms overexpressing the fully functional full length SMOC-1 (Figures 1,3C). Overexpressed SMOC-1(EC) also robustly rescued the Susm phenotypes of *smoc-1(0)* worms (Figure 3D). In contrast, overexpression of SMOC-1(TY) failed to rescue the *smoc-1(0)* body size (Figure 3C), and only slightly rescued the Susm phenotype of *smoc-1(0)* mutants (Figure 3D). The lack of rescue by the SMOC-1(TY) domain is not due to failed expression of SMOC-1(TY). As shown in Figure 3B, both SMOC-1(EC)::2xFLAG and SMOC-1(TY)::2xFLAG were detectable on western blots in worms overexpressing each corresponding domain. Taken together, our results indicate that the EC domain of SMOC-1 is both necessary and sufficient to regulate BMP signaling when overexpressed.

**Figure 3.**
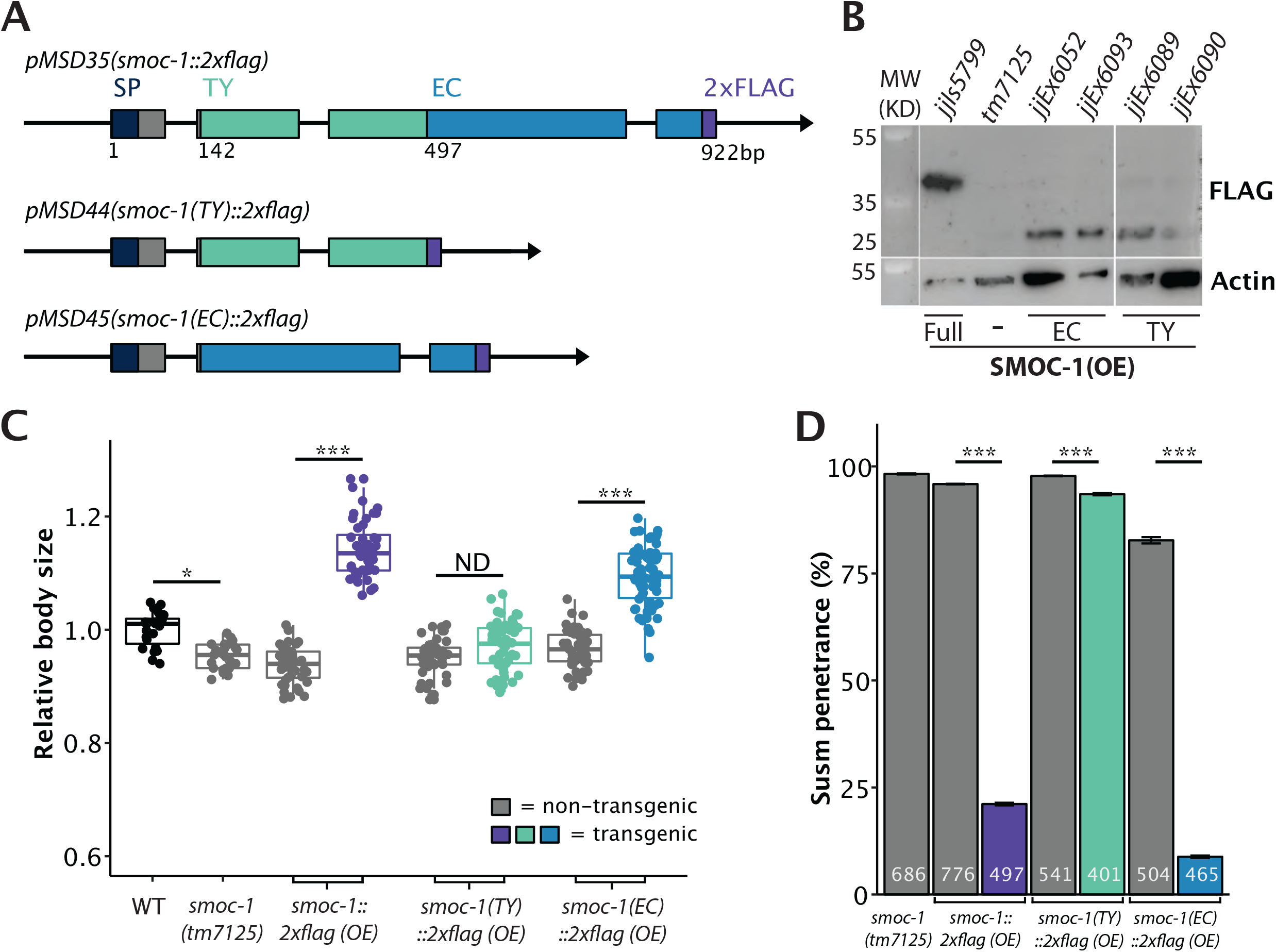
The EC domain of SMOC-1 is sufficient to regulate BMP signaling when overexpressed. **A)** Diagrams depicting the genomic constructs expressing full length SMOC-1::2xFLAG (pMSD35), SMOC-1(TY)::2xFLAG (pMSD44), and SMOC-1(EC)::2xFLAG (pMSD45). All plasmids contain the same 2kb promoter and 2kb 3’UTR of *smoc-1*. SMOC-1(TY) ends at amino acid 134, while SMOC-1(EC) starts at amino acid 135, both containing the same SMOC-1 signal peptide (SP), and the 2xFLAG tag. **B)** Western blot of 50 gravid adults of each indicated genotype, probed with anti-FLAG (top) and anti-actin antibodies (bottom). The strain overexpressing full length SMOC-1::2xFLAG (*jjIs5799*) is an integrated transgenic strain, while those overexpressing SMOC-1(TY)::2xFLAG (*jjEx6089* and *jjEx6090*) or SMOC-1(EC)::2xFLAG (*jjEx6052* and *jjEx6093*) carry the transgenes as extra chromosomal arrays, thus the expression level appeared lower due to random loss of the array during each mitotic division. **C)** Relative body sizes of strains carrying indicated versions of *smoc-1* as extra chromosomal arrays in a *smoc-1(tm7125)* null background at the same developmental stage (WT set to 1.0). For panels C and D, grey indicates non-transgenic worms that do not express any *smoc-1*. Two independent transgenic lines were measured and combined for each plasmid being tested here. Statistical analysis was done by comparing transgenic strains with non-transgenic counterparts. \*\*\**P*<0.001; \**P*<0.01, ND: no difference (ANOVA followed by Tukey HSD). WT: N=23. *tm7125*: N=25. Full length *smoc-1* (transgenic: N=44; non-transgenic: N=41). *smoc-1(TY)* (transgenic: N=40; non-transgenic: N=39). *smoc-1(EC)* (transgenic: N=66; non-transgenic: N=47). **D)** Summary of the Susm penetrance of strains carrying indicated versions of *smoc-1* in a *smoc-1(tm7125); sma-9(cc604)* background. The Susm penetrance refers to the percent of animals with one or two M-derived CCs as scored using the *arIs37(secreted CC::GFP)* reporter. For each genotype, two independent isolates were generated (as shown in the strain list), the Susm data from the two isolates were combined and presented here. Number of animals scored are noted on each bar. Statistical analysis was done to compare transgenic strains with non-transgenic counterparts. ***P<0.001 (general linear model, Wald statistic).

### The SMOC-1 EC domain is not fully functional when expressed at the *smoc-1* endogenous locus

To determine if SMOC-1(EC) is sufficient to function in BMP signaling when expressed at the endogenous level, we used CRISPR to alter the *smoc-1* genomic locus and generated an allele (*jj441)* that expresses SMOC-1(EC), and two alleles (*jj411* and *jj412*) that express SMOC-1(TY) (Figure 4A). We then tested the functionality of SMOC-1(EC) and SMOC-1(TY) using both the body size assay and the more sensitive Susm assay. We found that while SMOC-1(EC)-expressing worms do not exhibit any body size defect, SMOC-1(TY)-expressing worms are even smaller than *smoc-1(0)* null mutant worms (Figure 4B). Furthermore, both SMOC-1(EC)-expressing worms and SMOC-1(TY)-expressing worms exhibited a partially penetrant Susm phenotype (Figure 4C). Since worms carrying the WT *smoc-1* locus do not display any Susm phenotype (Figure 4C), our results suggest that both the TY and the EC domains are required for full function of SMOC-1 when expressed at the endogenous level.

**Figure 4.**
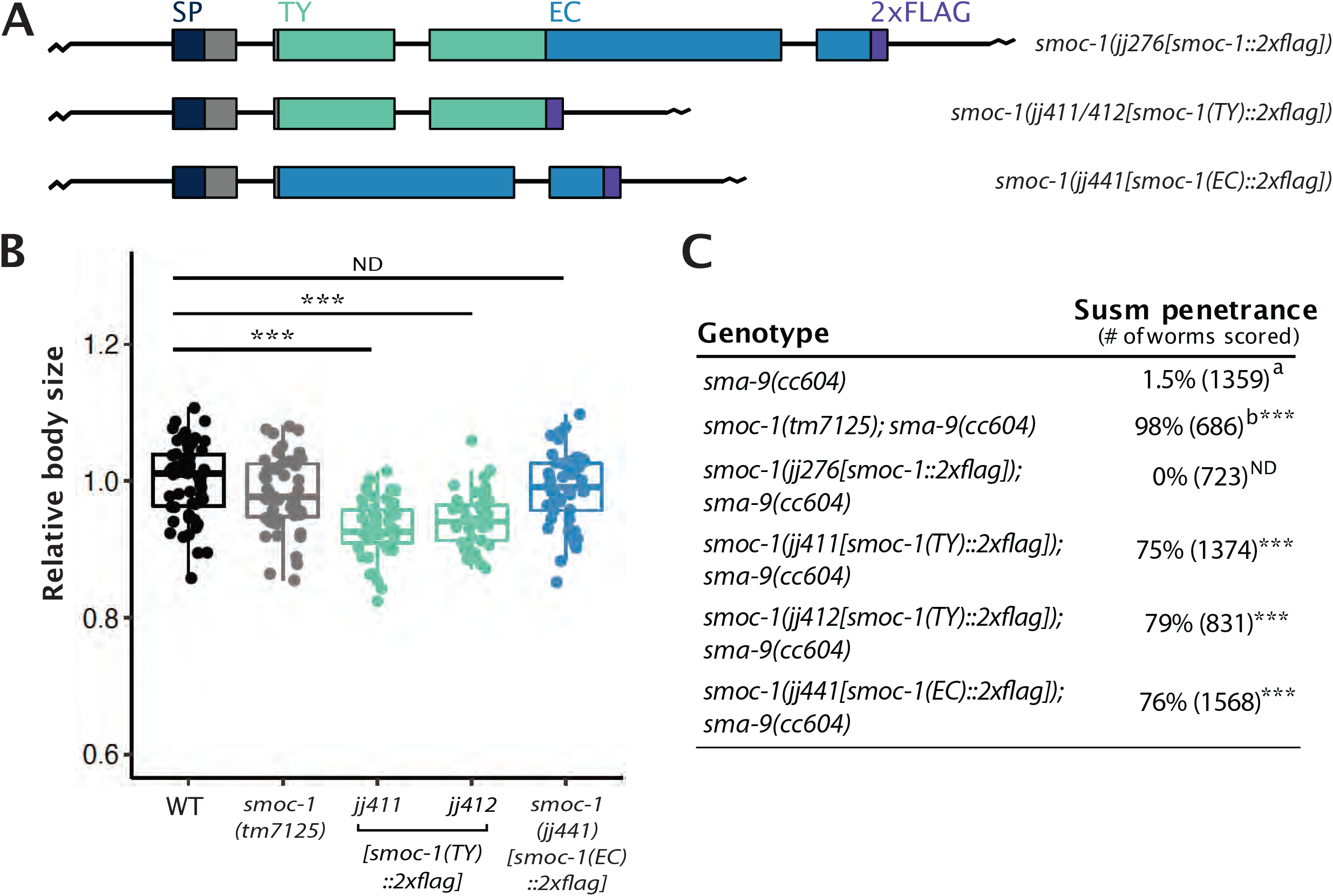
When expressed at the endogenous locus, neither SMOC-1(TY) nor SMOC-1(EC) is fully functional in regulating BMP signaling. **A)** Diagrams depicting the full length *smoc-1*, as well as truncated *smoc-1(TY)* and *smoc-1(EC)* at the endogenous locus. All of them are tagged with a 2xFLAG tag at the C-terminal end. **B)** Relative body sizes of worms at the same developmental stage WT (WT set to 1.0). \*\*\**P*<0.001; ND: no difference (ANOVA followed by Tukey HSD). WT: N=54. *tm7125:* N=51. *jj441*: N=51. *jj411*: N=47. *jj412*: N=41. **C)** Table showing the penetrance of the Susm phenotype of *smoc-1(TY)* and *smoc-1(EC)* as compared to the *smoc-1(0)* worms. ^a^ The lack of M-derived CCs phenotype is not fully penetrant in *sma-9(cc604)* mutants. ^b^ Data from DeGroot et al. [12]. Statistical analysis was conducted by comparing double mutant lines with the *sma-9(cc604)* single mutants. \*\*\**P*<0.001; ND: no difference (unpaired two-tailed Student’s *t*-test).

### The BMP ligand DBL-1 binds to full length SMOC-1, but not LON-2/glypican, *in vitro*

Previous studies of the *Drosophila* and Xenopus SMOC proteins led to a model where SMOCs function by competing with BMP ligands to bind to HSPGs, allowing the spreading of BMP ligands [17]. We have found that SMOC-1 can associate with LON-2/glypican via its EC domain, just like in these other systems. We therefore decided to further investigate the relationship between SMOC-1, the BMP ligand DBL-1, and LON-2/glypican, by expressing differently tagged SMOC-1, DBL-1 and LON-2 proteins using the S2 cell expression system (Figure 5). Since BMP molecules are produced as inactive molecules with a prodomain attached to the mature active domain ([28]), we generated constructs that would allow us to detect both the prodomain and the mature domain of DBL-1 (Figure 5A,C). We then performed reciprocal co-IP experiments for each protein pair. We did not detect any association between LON-2/glypican with either full length DBL-1 (Figure S2), or each of the domains of DBL-1 (Figure 5B). Instead, full length SMOC-1, but not SMOC-1(EC) nor SMOC-1(TY), can co-immunoprecipitate with the mature domain of DBL-1 (Figure 5D).

**Figure 5.**
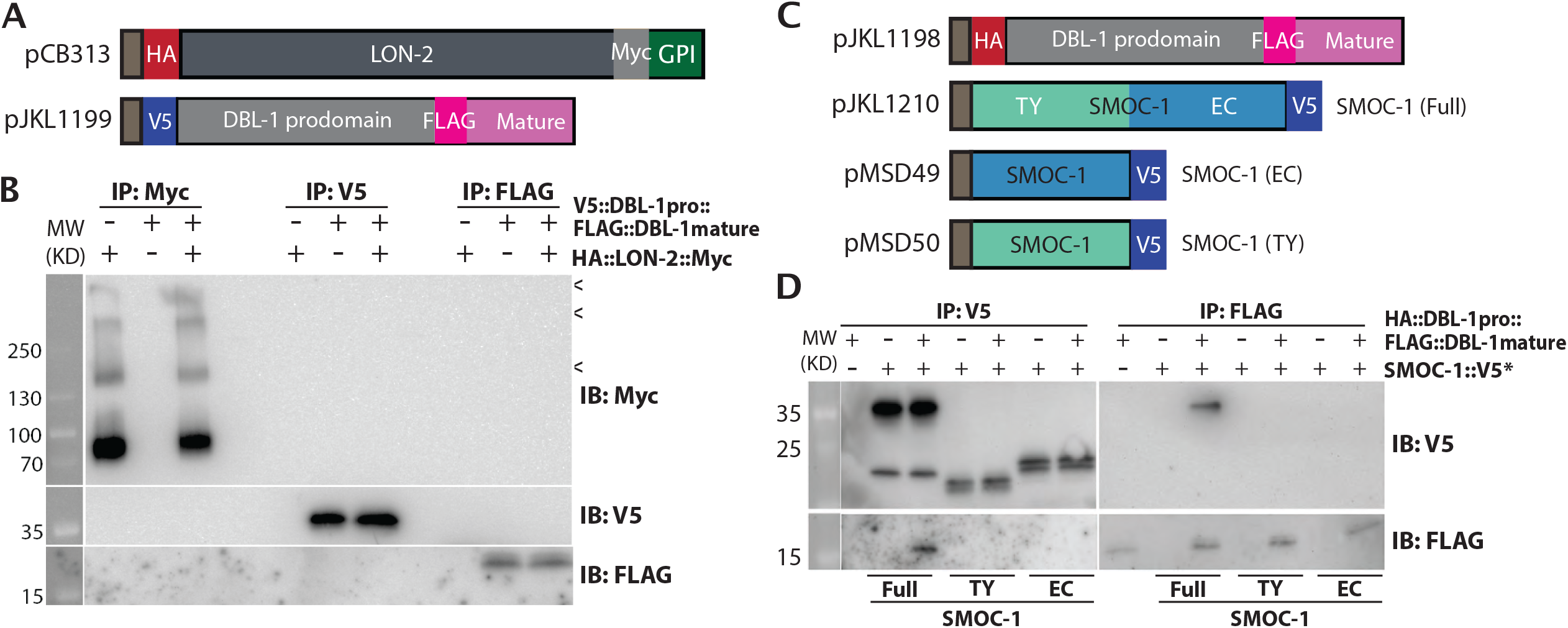
SMOC-1, but not LON-2, binds to DBL-1 when expressed in S2 cells. **A)** Diagrams of LON-2 and DBL-1 expression constructs used in the *Drosophila* S2 cell expression system. **B)** Results of co-IP experiments testing the interaction between HA::LON-2::Myc and V5::DBL-1 prodomain::FLAG::DBL-1 mature domain. Immunoprecipitation (IP) with anti-Myc beads, anti-V5 beads or anti-FLAG beads and immunoblot (IB) with anti-Myc, anti-V5 or anti-FLAG antibodies, as indicated. Experiments were independently repeated in triplicate, with representative results shown in this figure. < points to faint bands that likely represent glycosylated LON-2. **C)** Diagrams of SMOC-1 and DBL-1 expression constructs used in the *Drosophila* S2 cell expression system. **D)** Results of co-IP experiments testing the interaction between HA::DBL-1 prodomain::FLAG::DBL-1 mature domain and different versions of SMOC-1::V5*. Immunoprecipitation (IP) with anti-V5 beads or anti-FLAG beads and immunoblot (IB) with anti-V5 or anti-FLAG antibodies, as indicated. The source of DBL-1 in these experiments was cell media, which does not contain full length DBL-1, but only HA-tagged prodomain and the FLAG-tagged mature domain. Experiments were independently repeated in triplicate, with representative results shown in this figure.

### *In silico* structural modeling supports the interaction between LON-2/glypican and SMOC-1, and between SMOC-1 and DBL-1/BMP

The interaction between SMOC-1 and the mature domain of DBL-1/BMP was unexpected, given previous studies of SMOC proteins in other systems. We therefore sought independent ways to verify this finding. Recent advances in protein structure prediction have enabled the structures of protein-protein interactions to be modeled *in silico* ([29-32]). These predicted structural models can serve as useful tools for interpreting functional results and for generating hypotheses that can be tested through further experimentation. We used the ColabFold [33] implementation of AlphaFold2 [29] to determine whether confident structural predictions could be generated for the interactions between LON-2, SMOC-1, and mature DBL-1. Consistent with results of our physical interaction experiments, this structural modeling predicted a strong interaction between LON-2 and SMOC-1, and their interaction involves the EC domain of SMOC-1 (Figure 6A). None of the possible forms of DBL-1 (mature domain, pro-domain, or full-length) were predicted to interact with LON-2. However, the mature domain of DBL-1 and SMOC-1 were predicted to interact, and the predicted interaction involves a bipartite interaction with both the TY domain and the C-terminal portion of the EC domain of the full-length SMOC-1 protein (Figure 6B). All these predictions are consistent with results of our physical interaction experiments.

**Figure 6.**
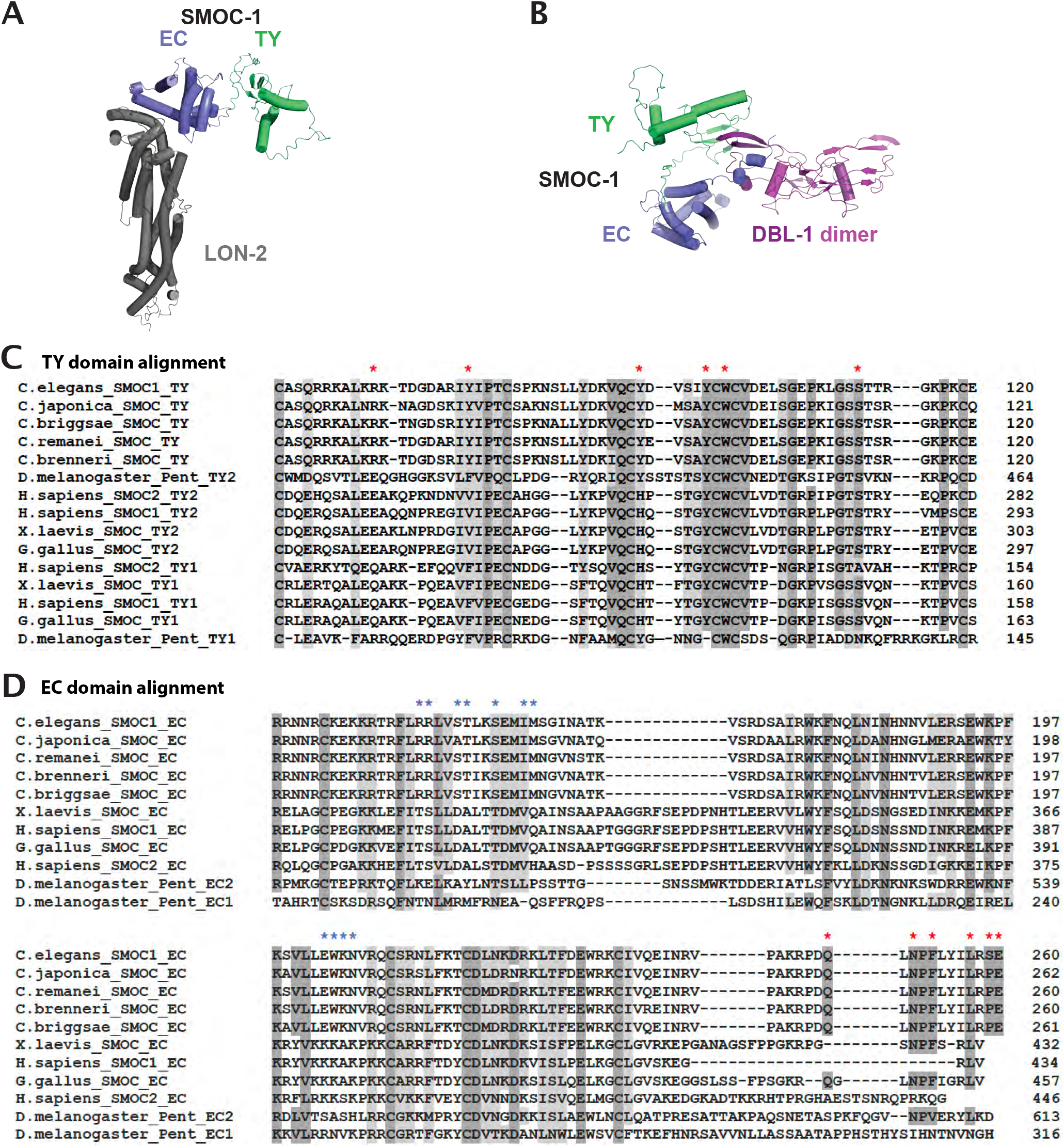
Structural modeling of interactions between SMOC-1 and LON-2, and between SMOC-1 and DBL-1. **A)** Predicted structure of a complex formed between LON-2 and SMOC-1. The EC domain of SMOC-1 is predicted to interact with LON-2. **B)** Predicted structure of a complex formed between SMOC-1 and a homodimer of the DBL-1 mature domain. Both the EC and TY domains of SMOC-1 are predicted to interact with DBL-1. **C**) Multiple sequence alignment of the TY domains of SMOC-1 homologs using Clustal Omega (CLUSTAL O(1.2.4)) Red * marks the residues at the interface between SMOC-1 and DBL-1, as identified via ColabFold. D) Multiple sequence alignment of the EC domains of SMOC-1 homologs using Clustal Omega (CLUSTAL O(1.2.4)) Red * marks the residues at the interface between SMOC-1 and DBL-1, and blue * marks the residues at the interface between SMOC-1 and LON-2, as identified via ColabFold. In both C and D, dark shaded residues are identical, while light shaded residues are conserved, among all or most of the homologs.

Importantly, our structural modeling also identified key residues located at the interfaces between each pair of interacting partners. Sequence comparisons between the *C. elegans* SMOC-1, LON-2, DBL-1 and their corresponding counterparts in other organisms, ranging from other nematode species to *Drosophila* and mammals, showed that these key residues are highly conserved among each of the three protein families (Figure 6C-D, S3, S4). In particular, a NPF/VxxxL motif at the C-terminal end of the EC domain appears to be conserved among SMOC homologs in *Drosophila, Xenopus*, and chicken.

### LON-2/glypican, SMOC-1 and DBL-1/BMP forms a tripartite complex *in vitro*

When we included all three proteins in the structural prediction together, a structure was predicted in which LON-2 interacts with SMOC-1 and SMOC-1 interacts with mature DBL-1 (Figure 7A-B). This prediction suggested the presence of a SMOC-1-dependent tripartite complex between LON-2/glypican, SMOC-1 and DBL-1/BMP.

**Figure 7.**
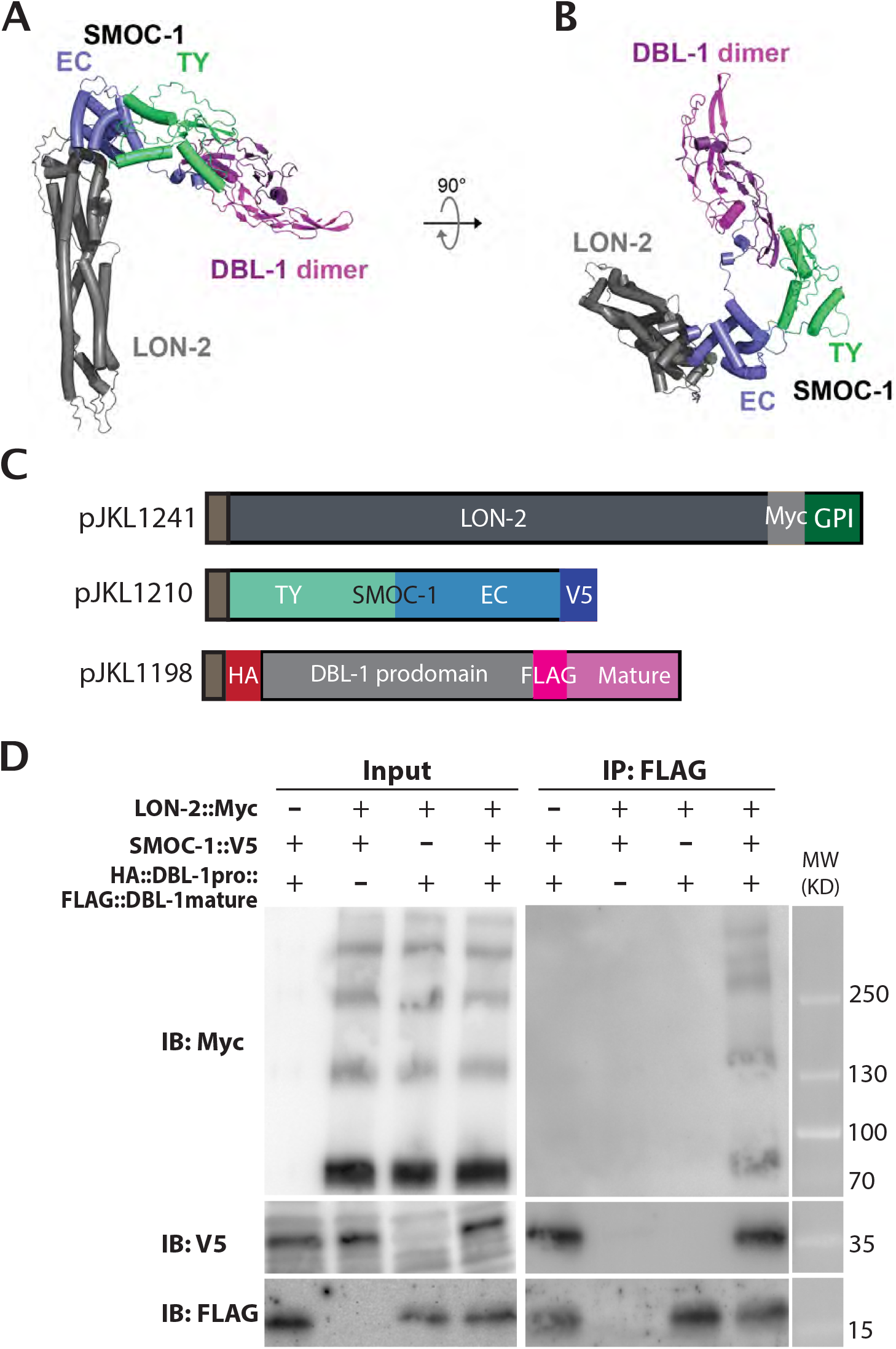
co-IP results testing the model (for tripartite complex formation) Predicted structure of a complex formed between LON-2, SMOC-1, and a homodimer of the DBL-1 mature domain. The membrane-anchoring region of LON-2 lies at the bottom of the panel. The same structure prediction as in A), but shown from the ‘top’. **C**) Diagrams of LON-2, SMOC-1 and DBL-1 expression constructs used in the *Drosophila* S2 cell expression system. **D)** Results of co-IP experiments testing the interaction between LON-2::Myc and HA::DBL-1 prodomain::FLAG::DBL-1 mature domain in the presence or absence of SMOC-1::V5. Immunoprecipitation (IP) with anti-FLAG beads and immunoblot (IB) with anti-Myc, anti-V5 or anti-FLAG antibodies, as indicated. Experiments were independently repeated in triplicate, with representative results shown in this figure.

We experimentally tested this *in silico* structural prediction, using proteins expressed in the *Drosophila* S2 expression system. We performed co-immunoprecipitation experiments by mixing LON-2 and DBL-1 together in the presence or absence of SMOC-1. Again, the mature domain of DBL-1 can co-IP SMOC-1, and this interaction is irrespective of whether LON-2 is present or not (Figure 7C). In the same co-IP experiments, IP using the mature domain of DBL-1 pulled down LON-2 only in the presence of SMOC-1 (Figure 7C). These results strongly suggest that SMOC-1 can mediate LON-2 and DBL-1 interaction, and that the three proteins can form a SMOC-1-dependent tripartite complex.

### A model for how SMOC-1 functions to regulate BMP signaling

LON-2/glypican is well established to be a negative regulator of BMP signaling in *C. elegans* [34]. We have previously shown that the other glypican in the *C. elegans* genome, GPN-1/glypican, does not function in BMP signaling [35]. In contrast to LON-2, SMOC-1 can positively promote BMP signaling when overexpressed ([12], and this study). Our co-IP results showed that SMOC-1 can bind to both LON-2 and the mature domain of DBL-1, and that it can mediate the formation of a LON-2-SMOC-1-DBL-1 tripartite complex. Based on these data, we proposed the following model for how SMOC-1 functions to regulate BMP signaling in *C. elegans* (Figure 8A). In our model, SMOC-1 has both a negative, LON-2-dependent role and a positive, LON-2-independent role in regulating BMP signaling. On the one hand, SMOC-1 binds to both the mature domain of DBL-1/BMP and LON-2/glypican, resulting in the sequestration of DBL-1/BMP, preventing DBL-1/BMP from interacting with the BMP receptors and inhibiting BMP signaling. On the other hand, secreted SMOC-1 can bind to the mature domain of DBL-1/BMP, possibly facilitating the movement of DBL-1/BMP through the extracellular space or the delivery of DBL-1/BMP to its receptors, thus promoting BMP signaling. The duality of SMOC-1 function in our model is consistent with previous findings that *smoc-1(0)* mutants are only slightly (∼5%) smaller than wildtype worms, and that *smoc-1(0); lon-2(0)* double mutants have an intermediate body size between *smoc-1(0)* and *lon-2(0)* mutants [12].

**Figure 8.**
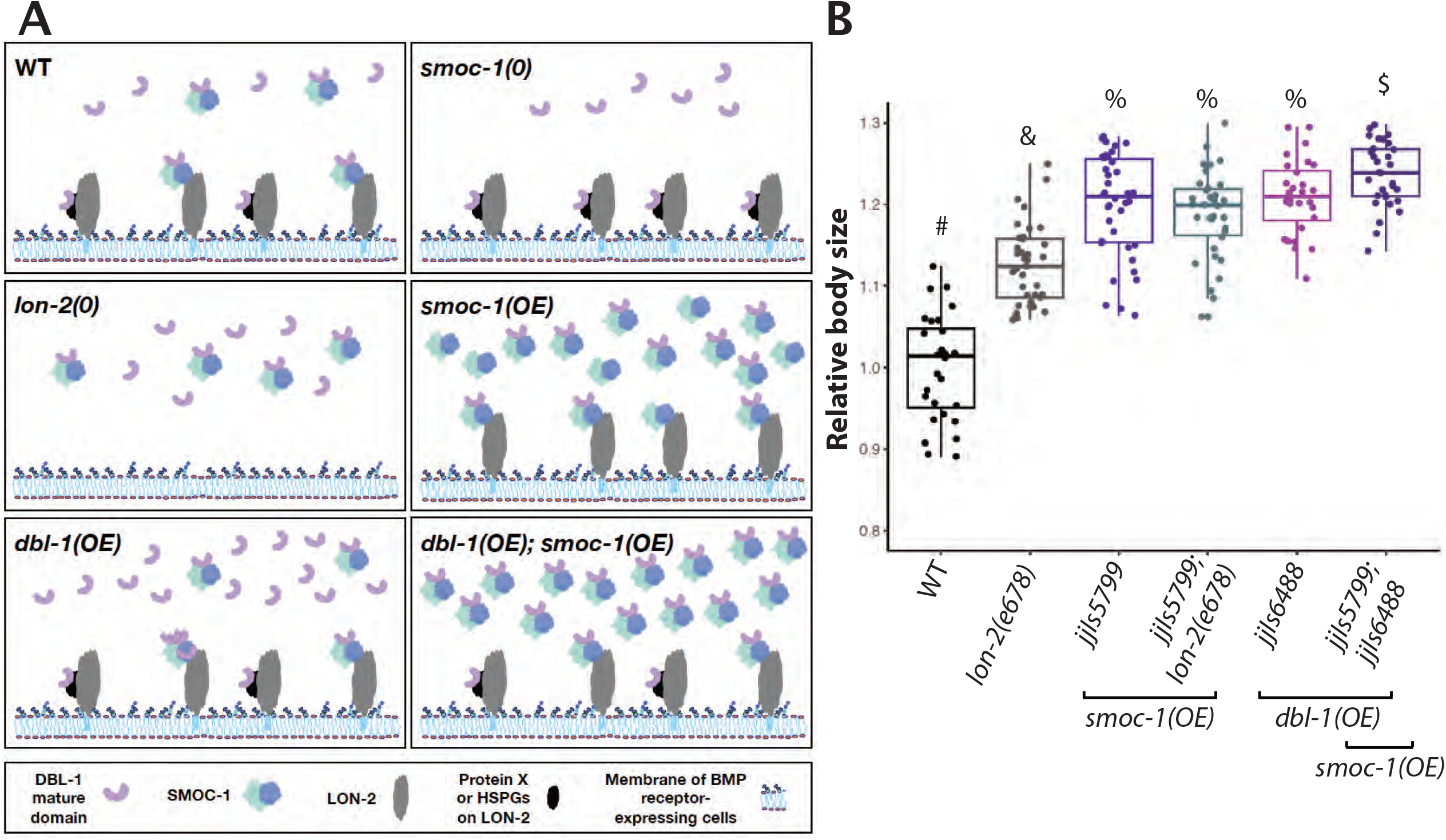
A model for how SMOC-1 functions to regulate BMP signaling. **A)** In wildtype (WT) animals, SMOC-1 is proposed to both negatively and positively regulate BMP signaling. For its negative role, SMOC-1 binds to LON-2/Glypican, which is expressed in cells that also express the BMP receptors (such as hypodermal and intestinal cells) to sequester DBL-1/BMP. In the meantime, SMOC-1 binds to DBL-1 and helps DBL-1 move through the extracellular environment, promoting BMP signaling. We propose that DBL-1, when bound to SMOC-1, either moves through the extracellular environment more efficiently or functions more effectively, than unbound DBL-1. At present, we cannot rule out the possibility that DBL-1 can also be sequestered by LON-2 either via another protein X or via the HSPGs on LON-2. Specific scenarios in different mutants are also depicted based on this model. **B**) Relative body sizes of various strains at the same developmental stage (WT set to 1.0). *lon-2(e678)* is a null allele of *lon-2. jjIs57799* is an integrated multicopy array line that overexpresses SMOC-1::V5. *jjIs6448* is an integrated multicopy array line that overexpresses DBL-1. Groups marked with distinct symbols are significantly different from each other (*P*<0.0001, in all cases when there is a significant difference), while groups with the same symbol are not. Tested using an ANOVA with a Tukey HSD. WT: N=28. *e678*: N=37. *jjIs5799:* N=39. *jjIs5799; e678*: N=34. *jjIs6448*: N=29. *jjIs5799; jjIs6448*: N=45.

According to our model, drastically overexpressed SMOC-1 can promote BMP signaling irrespective of the presence or absence of LON-2. Moreover, overexpressed SMOC-1 would be expected to further augment the positive effect of increased levels of DBL-1/BMP on BMP signaling. To test these hypotheses, we generated strains that overexpress *smoc-1 (jjIs5799[smoc-1(OE)])*, or *dbl-1 (jjIs6448[dbl-1(OE)])* (Figure S5) and measured the body sizes of animals overexpressing *smoc-1* in the presence or absence of wild-type *lon-2*, and in the presence or absence of *dbl-1(OE)*. As shown in Figure 8B, *smoc-1(OE)* animals are significantly longer than *lon-2(e678)* null animals. Notably, *smoc-1(OE); lon-2(e678)* double mutant animals are similar in length to *smoc-1(OE)* animals yet longer than *lon-2(e678)* null animals, demonstrating that overexpression of SMOC-1 can in fact promote BMP signaling in the absence of LON-2. Double mutants that overexpress both SMOC-1 and DBL-1 are significantly longer than either *smoc-1(OE)* or *dbl-1(OE)* single mutants (Figure 8B). Taken together, these genetic findings strongly support our model on the dual functions of SMOC-1 in regulating BMP signaling.

## DISCUSSION

In this study, we discovered that *C. elegans* SMOC-1 binds the glypican LON-2, as well as the mature domain of the BMP protein DBL-1. Moreover, SMOC-1 can mediate the formation of a glypican-SMOC-BMP tripartite complex. Our biochemical and molecular genetic data suggested dual functionality of SMOC-1, acting both positively and negatively in the BMP pathway.

The concept of SMOCs acting as both antagonists and expanders of BMP signaling has been proposed by Thomas and colleagues [17]. Xenopus XSMOC-1 and the *Drosophila* SMOC homolog Pent have been shown to promote the spreading of the BMP ligand, thus expanding the range of BMP signaling, both *in vitro* and *in vivo* [10, 11, 17, 36]. Both XSMOC-1 and human SMOC-1, can bind to heparin and heparan sulfate (HS), and in the case of *Drosophila* Pent, the BMP co-receptors Dally/glypican and Dally-like/glypican [10, 17, 26, 27]. Studies in *Drosophila* further showed that the ability of Pent to extend the range of BMP signaling is dependent on Dally/glypican in wing imaginal discs [10]. Because neither XSMOC-1 nor Pent have been found to bind BMPs, but BMPs can bind HSPGs [10, 17, 18, 27], a model was proposed where the binding of SMOCs to HSPGs competitively reduces the interaction between HSPGs and BMPs, thus promoting the spreading of BMPs [17].

At present, there is no unified model on how SMOCs function to inhibit BMP signaling. Using a Xenopus animal cap assay, Thomas and colleagues [16, 17] showed that both XSMOC-1 and Pent can inhibit BMP signaling downstream of the BMP receptor, at least for XSMOC-1, by activating the MAPK pathway. In contrast, *Drosophila* Pent has been shown to function upstream of receptor activation to inhibit BMP signaling in zebrafish [10]. A more recent study showed that mouse SMOC2 can inhibit BMP signaling by competitively binding to BMPR1B [13]. While our work supports the dual functions of SMOCs in regulating BMP signaling, it suggests a different model on how SMOC-1 can accomplish these roles in *C. elegans*. Our results are consistent with SMOC-1 functioning in a LON-2-dependent manner to negatively regulate BMP signaling, and in a LON-2-independent, but BMP-dependent, manner to promote BMP signaling.

### Glycosaminoglycan (GAG)-dependent and -independent interactions between SMOC-1 and LON-2/glypican

Using both IP-MS of worm extracts and co-immunoprecipitation of proteins expressed in *Drosophila* S2 cells, we have found that SMOC-1, specifically the EC domain of SMOC-1, can bind to LON-2/glypican. Our findings are consistent with previous findings showing that *Drosophila* Pent can co-IP with Dally/glypican and Dally-like/glypican, while XSMOC-1, hSMOC2 and Pent can all bind to heparin and heparan sulfate (HS) *in vitro* [10, 17, 26, 27]. The EC domains of SMOCs all have a stretch of positively charged residues that was hypothesized to mediate its interaction with HSPGs [26]. Fascinatingly, our protein structural modeling suggests that the SMOC EC domain may directly interact with the protein core of HSPGs. As shown in Figure 7, the key residues identified to mediate SMOC-1-LON-2 interaction are conserved among nematode SMOC proteins, as well as vertebrate SMOCs. Further, our results in *C. elegans* provide additional clarity to some previously perplexing findings by Taneja-Bageshwar and Gumienny [37, 38]. These authors showed that overexpressing the LON-2/glypican protein core is sufficient to rescue the long (Lon) body size phenotype of *lon-2(0)* null mutants, suggesting that the LON-2/glypican protein core is sufficient to inhibit BMP signaling. Their previous findings further showed that the N-terminal region of LON-2, LON-2(1-368), is sufficient to inhibit BMP signaling. Notably, this region contains all the key residues in LON-2 that mediate LON-2-SMOC-1 interaction, as identified by our structural modeling (between amino acids 100 and 325, Figure S3). We therefore postulate that part of LON-2’s function in regulating BMP signaling is mediated by the interaction of SMOC-1 with the LON-2 protein core. Similarly, Kirkpatrick and colleagues found in *Drosophila* that the protein core of Dally/glypican is partially functional, while its overexpression actually limits Dpp/BMP signaling in the wing imaginal discs [39]. Whether Pent is involved in mediating this function of Dally/glypican is unclear. In the *C. elegans* study described above, Taneja-Bageshwar and Gumienny [37, 38] also showed that the heparan sulfate (HS) attachment sites are important for a truncated form of LON-2 (aa423-508) to function in inhibiting BMP signaling. It is therefore still possible that SMOC-1, like its homologs in other systems, can also bind to LON-2 via the polysaccharide heparan sulfate chains on LON-2. Future research will be needed to identify the key signatures in LON-2 that mediate its interaction with SMOC-1.

### SMOC-1 as a mediator of glypican and BMP interaction

LON-2/glypican is a well-established negative regulator of BMP signaling in *C. elegans* [34]. Previous studies showed that when expressed in mammalian HEK293T cells, membrane-tethered LON-2 can bind to mammalian BMP2 after chemical crosslinking [34]. Since *Drosophila* and vertebrates BMPs have been found to bind to HSPGs [18, 40], a model was proposed where LON-2/glypican negatively regulates BMP signaling by sequestering the DBL-1/BMP ligand in *C. elegans* [34]. We did not detect any interaction between LON-2 and any form of DBL-1 (Figures 5, S2). Although we cannot rule out the possibility that post-translational modifications happening in *Drosophila* S2 cells are different from those found in *C. elegans*, we believe that the lack of LON-2 and DBL-1 interaction in our S2 cell expression system is unlikely due to the lack of glycosaminoglycan (GAG) modifications of LON-2. As shown in Figures 2, 5, and S2, we detected multiple higher molecular weight bands on western blots that likely correspond to GAG-decorated LON-2.

A plausible model to reconcile previous data [34] and ours is that LON-2 may bind to DBL-1 indirectly via a bridging molecule. Based on our findings, we believe that SMOC-1 is the bridging molecule. We detected strong interaction between SMOC-1 and the mature domain of DBL-1 (Figure 5). Furthermore, we found that SMOC-1 can mediate the formation of a tripartite complex between LON-2 and DBL-1 (Figure 7). *In silico* structural modeling using the ColabFold [33] implementation of AlphaFold2 [29] supported the interactions between LON-2, SMOC-1, and mature DBL-1 (Figures 6, 7). Importantly, key amino acids identified by structural modeling in each of the three proteins to mediate pair-wise protein-protein interactions appear to be highly conserved (Figures 6, S3, S4). Our data, combined with the previous model proposed by Gumienny and colleagues [34], suggest that LON-2/glypican acts as a negative regulator of BMP signaling in *C. elegans* by sequestering the DBL-1/BMP ligand via SMOC-1, and that SMOC-1 facilitates the negative regulation of BMP signaling by forming a tripartite complex with LON-2/glypican and DBL-1/BMP. Since SMOC homologs from *C. elegans, Drosophila*, and Xenopus all functionally interact with HSPGs, we propose that the SMOC-HSPG axis is an evolutionarily conserved module important in regulating BMP signaling.

Our previous work [12] and this work showed that when overexpressed, SMOC-1 can act as a positive regulator of BMP signaling. We argue that this positive role of SMOC-1 is independent of LON-2/glypican. We have previously shown that *smoc-1(0); lon-2(0)* double null mutants exhibit an intermediate body size phenotype when compared to each single mutant [12]. In this study, we showed that overexpressing SMOC-1 either in the wild-type background or in the *lon-2(0)* null background produced a similar yet longer body size phenotype than *lon-2(0)* single null mutants (Figure 8B), suggesting that SMOC-1 can promote BMP signaling beyond counteracting the negative effects of LON-2. It is unlikely that a LON-2 paralog is mediating this function, as we have previously shown that GPN-1, the other glypican in the *C. elegans* genome, does not play a redundant role with LON-2 in regulating BMP signaling [35]. We have previously shown that the positive role of SMOC-1 in regulating BMP signaling is dependent on the presence of the BMP ligand [12]. Since we detected strong interaction between SMOC-1 and DBL-1/BMP, we argue that by binding to DBL-1/BMP, SMOC-1 facilitates either the movement of DBL-1/BMP through the extracellular space or the binding of DBL-1/BMP to its receptors, thus promoting BMP signaling. Consistent with this notion, strains overexpressing both SMOC-1 and DBL-1 exhibit even a longer body size than strains overexpressing either SMOC-1 or DBL-1 alone, or *lon-2(0)* null mutants (Figure 8B). Collectively our results strongly suggest that the positive role of SMOC-1 in regulating BMP signaling is independent of LON-2/glypican. Whether SMOC-1 accomplishes this role alone or with the help of another protein(s) is currently unknown.

### The importance of both the TY and the EC domains for SMOC-1 function

We have shown that both the TY and the EC domains are required for full function of SMOC-1 at the endogenous locus. The requirement for both the TY and the EC domains for full SMOC-1 function is consistent with both of our co-IP and structural modeling results, which demonstrated that residues in both the TY and the EC domains are involved in the interaction between SMOC-1 and DBL-1 (Figures 6 and S4). However, paradoxically, overexpression of SMOC-1(EC) is sufficient to promote BMP signaling. One possible explanation for this minor inconsistency is that under normal expression levels, the interaction between SMOC-1 and mature DBL-1 requires both the TY and EC domains, but when overexpressed the EC domain is sufficient for interaction with DBL-1. Consistent with this notion, we have previously shown that two single amino acid substitution mutations in the respective TY domain and EC domain of SMOC-1, *jj85(E105K)* and *jj65(C210Y)*, display partial loss-of-function phenotypes [12]. Yet, like overexpressing SMOC-1(EC), overexpressing either SMOC-1(E105K) or SMOC-1(C210Y) was also sufficient to promote BMP signaling (Figure S5). Since many functional assays in *C. elegans* involve the utilization of repetitive transgenic arrays that can cause overexpression of the transgene, our results also highlight the importance of carrying out functional studies of proteins at the endogenous expression levels.

Our work and previous studies on SMOC-1 homologs have shown that the EC domain of SMOC-1 can bind to HSPGs. What is the function of the TY domain in SMOC-1? The thyroglobulin type-1 (TY) domain is evolutionarily conserved across metazoans and seen in a variety of functionally diverse proteins, including thyroglobulin, SMOCs, nidogens, and IGFBPs [41, 42]. The TY (type 1a) domain is characterized by six conserved cysteine residues, including ‘QC’ and ‘CWCV’ residue motifs, that form intramolecular disulfide bridges [43]. Outside of these conserved regions, the TY domain has highly variable loops which may contribute to the wide variety of associated functions. TY domain-containing proteins may function to inhibit cysteine cathepsins, while others inhibit aspartate peptidases, papain-like proteases, or metalloproteases [41, 42]. Testicans, which contain a TY domain as well as an EC domain, can work as a competitive inhibitor of cathepsin L as found in human cell culture. On the other hand, human testicans have also been found to bind to cathepsin L to act as a chaperone, preventing degradation and encouraging cathepsin activity within the ECM [44, 45]. Thomas and colleagues found that one of the TY domains (TY1) in XSMOC-1 is required for XSMOC-1’s function in inhibiting BMP signaling [17]. How it accomplishes this function, however, is not understood. Future research will determine how the SMOC-1 TY domain functions and whether it acts in any analogous ways as other TY-containing proteins.

## Conclusion

There are two SMOC homologs in mammals, SMOC1 and SMOC2. Both mice and human individuals with mutations that severely reduce SMOC-1 expression exhibit abnormal tooth, eye, and limb development [14, 15, 46-48]. Mutations in hSMOC2 have been found to be associated with defects in tooth development [49, 50] and vitiligo [14, 51, 52]. Abnormal expression of SMOCs has also been associated with multiple cancers as well as kidney and pulmonary fibrosis [53-59]. We have previously shown that both hSMOC1 and hSMOC2 can partially rescue the BMP signaling defect of *smoc-1(0)* null mutants in *C. elegans* [12]. Therefore, future research on the mechanistic details of SMOC function to regulate BMP signaling in an *in vivo* system such as *C. elegans* may have significant implications for human health.

## MATERIALS AND METHODS

### Plasmid constructs and transgenic lines

All plasmids and oligonucleotides used in this study are listed in Tables S2 and S3, respectively. Plasmids used for expression in *Drosophila* S2 cells were generated from pCB313 (gift from Dr. Claire Bénard, [60]).

Transgenic strains were generated using pCFJ90[*myo-2p::mCherry::unc-54 3’UTR*] (gift from Erik Jorgensen) or LiuFD290[*ttx-3p::mCherry*] (gift from Oliver Hobert) as co-injection markers. Two transgenic lines with the best transmission efficiency across multiple generations were analyzed with each plasmid. Integrated transgenic lines were generated using gamma irradiation, and then outcrossed with wildtype N2 worms at least two times. Table S4 lists all the strains generated in this study.

For IP/MS experiments, integrated transgenic lines overexpressing untagged SMOC-1 or SMOC-1::2xFLAG were used (see Table S4). Transgenic strains overexpressing either SMOC-1(TY)::2xFLAG or SMOC-1(EC)::2xFLAG were generated using an embryonic lethal temperature-sensitive *pha-1(e2123)* background, by including a *pha-1* expressing plasmid (pC1) and a florescent co-injection marker within the array. Only transgenic worms are viable at the restrictive temperature (25°C) [61]. The florescent co-injection marker allowed for visual assessment of multi-generational transgene transmission efficiency. The usage of these two markers allowed us to grow large populations of exclusively transgenic worms carrying the transgenes at high transmission efficiency.

### Generating endogenous smoc-1::2xflag, smoc-1(TY)::2xflag, smoc-1(EC)::2xflag and HA::dbl-1 strains using CRISPR/Cas9-mediated homologous recombination

All sgRNA guide plasmids were generated using the strategy described in[62, 63]. To generate each specific modifications, the specific sgRNA plasmids (see Table S2) were injected with the Cas9-encoding plasmid, pDD162 (Dickinson, Pani et al. 2015), and the single-strand oligodeoxynucleotide homologous repair template, into Wildtype N2 worms. pRF4(*rol-6(d)*) was used as a co-injection marker. Injected P0 animals were singled. F1 progeny were picked from plates that gave the most roller progeny, allowed to lay eggs and screened for successful CRISPR events via PCR (see Table S3 for oligo information). The resulting stable strains were confirmed by genotyping and Sanger sequencing. The *smoc-1(TY)::2xflag* and the *smoc-1(EC)::2xflag* strains were generated in the *smoc-1(jj276[smoc-1::2xflag])* background using either single-strand oligodeoxynucleotide or plasmid as homologous repair template (see Tables S2 and S3).

### Generating strains containing integrated extra-chromosomal arrays overexpressing either smoc-1::2xflag or HA::dbl-1

Genomic DNA including the coding region of *smoc-1::2xflag*, 2kb upstream and 2kb downstream sequences was amplified from *jj276* worms using JKL1549 and JKL1550 and cloned into a pBSII SK+ vector to generate the plasmid pMSD35. Transgenic strains were generated using *ttx-3p::mCherry* (LiuFD290) as a co-injection marker. Integrated transgenic lines (*jjIs5798, jjIs5799, jjIs5800*) were generated using gamma-irradiation, followed by two rounds of outcrossing with N2 worms. Similar approaches were used to generate pTYC3, which has sequences corresponding to the genomic DNA that includes the coding region of *HA::dbl-1*, 3kb upstream and 0.7kb downstream sequences, cloned into a pBSII SK+ vector. Transgenic strains were generated using pCFJ90[*myo-2p::mCherry::unc-54 3’UTR*] as a co-injection marker. A spontaneously integrated strain (*jjIs6448*) was generated, and then outcrossed for three rounds with wildtype worms.

### Body size measurements

Body size measurements were conducted as previously described [12, 64]. Gravid adults were bleach synchronized, with resulting embryos incubated in M9 buffer rotating at 15°C until hatched (24-48h). Synchronized L1s were plated and grown at 20°C until the L4.3 vulval stage was seen in a majority of worms. For imaging, worms were washed from plates, treated with 0.3% sodium azide until straightened, and then mounted onto 2% agarose pads. Images were taken using a Hamamatsu Orca-ER camera using the iVision software (Biovision Technology). Using Fiji, worm body lengths were measured from images using the segmented line tool. An ANOVA and Tukey’s honest significant difference (HSD) was used to test for differences in body size between genotypes using R (https://www.R-project.org/).

### Suppression of sma-9(0) M-lineage defect (Susm) assay

For the suppression of *sma-9(0)* M-lineage defect (Susm) assay, worms were grown at 20°C, and the number of adult animals with four coelomocytes (CCs) and five-to-six CCs were tallied across multiple plates [65]. The reported Susm penetrance refers to the percent of animals with one or two M-derived CCs as scored using the CC::GFP reporter. For each genotype, two independent isolates were generated (as shown in the strain list in Table S3), three to seven plates of worms from each isolate were scored for the Susm phenotype, and the Susm data from the two isolates were combined and presented. The lack of M-derived CCs phenotype is not fully penetrant in *sma-9(cc604)* mutants [28]. For the Susm rescue experiments, we used R to generate a general linear model with binomial error and a logit link function designating transgenic state as the explanatory function. The Wald statistic test was used to determine if transgenic state (transgenic vs. non-transgenic worms within the same line) is associated with CC number.

### Preparation of worm lysates for SDS-PAGE and western blot analysis

Western blot of *C. elegans* was conducted to detect proteins expressed *in vivo*. Indicated number of worms were picked into 20ul of double distilled water, and lysed by addition of 5ul of 5xSDS buffer (0.2 M Tris⋅HCl, pH 6.8, 20% glycerol, 10% SDS, 0.25% bromophenol blue, 10% β-mercaptoethanol) followed by snap freezing in liquid nitrogen. Samples were heated to 95°C for 10 minutes, stored at -20°C, and used for subsequent SDS-PAGE and western blot analysis.

### Immunoprecipitation of FLAG tagged proteins from C. elegans

*C. elegans* strains were repeatedly bleach synchronized and grown on 90mm NGM plates seeded with NA22 bacteria, until desired population size was reached. Approximately 25 NA22 plates containing about 10,000 synchronized gravid adults were bleached to get a target population of about 2,000,000 or more synchronized L1s in M9. Approximately 4,000,000 synchronized L1s were plated on 15cm egg plates (NGM strep with OP50-1 and chicken egg; [66] and grown until population reached the L4 stage (48 hours at 25°C). Worms were washed from plates and collected with H150 (50mM HEPES pH 7.6, 150mM KCl). Successive pelleting and washing were done to remove any excess food or debris. Finally, worms were pelleted and an equal volume of H150g10 (H150 with 10% glycerol) with protease inhibitor (Pierce, A32965) was added to make a worm slurry. “Worm popcorn” was made by adding the slurry dropwise directly into liquid nitrogen. Resulting popcorn was stored in 50mL conical tubes at -80°C.

To physically break worms, a mortar and pestle was used to grind the popcorn until no-to-few intact worms were visible. For the first IP/MS experiment, 20g of popcorn was used per strain, while 10g of popcorn was used in the second IP/MS experiment. Samples were kept in liquid nitrogen throughout the process to avoid unwanted thawing. Worm homogenates were then thawed on ice and diluted 1:5 (w:v) with H150g10 with 1% Triton. Samples were centrifuged at 12,000g at 4°C to separate soluble and insoluble fractions. Soluble fraction was filtered using a 0.45µm filter (Fisher brand Disposable PES Bottle Top filters, FB12566511) before being added to pre-equilibrated anti-Flag M2 magnetic beads (Millipore Sigma, M8823) for incubation overnight at 4°C with rotation. After incubation, unbound fraction was removed and beads were washed three times with H150g10 to remove unbound proteins. A final wash in TBS (20mM Tris HCl, 150mM NaCl, pH 7.6) was done before eluting with FLAG peptide (Sigma, F3290). Elution was done by adding 5x volumes of packed bead volume of FLAG peptide in TBS at 150ng/µl, followed by incubation at 4°C for 30mins. This was repeated for each sample and the two eluates were pooled together.

### Mass spectrometry

Eluates were submitted to Biotechnology Resource Center (BRC) at Cornell University for analysis by mass spectrometry (MS). Briefly, samples were prepared by in-solution trypsin digestion before conducting nanoLC-ESI-MS/MS analysis using an Orbitrap Fusion™ Tribrid™ (Thermo-Fisher Scientific, San Jose, CA) mass spectrometer equipped with a nanospray Flex Ion Source, and coupled with a Dionex UltiMate 3000 RSLCnano system (Thermo, Sunnyvale, CA) [67, 68]. Processing workflow used SequestHT and MS Amanda with Percolator validation. Database search was conducted against a *Caenorhabditis elegans* database downloaded from NCBI in June 2021. Only high confidence peptides defined by Sequest HT with a 1% FDR by Percolator were considered for confident peptide identification. Abundance ratios relative to untagged (eg. *smoc-1::2xflag/smoc-1*) were assessed to identify candidate interaction partners, with values over 2.0 being considered enriched. Only hits with two or more mapped peptides were considered here.

### Coimmunoprecipitation of proteins expressed in Drosophila S2 cells

*Drosophila* S2 cells were grown in M3+BPYE+10% Heat-inactivated FBS and transfected using a calcium phosphate method (Invitrogen protocol, Version F 050202 28-0172). For non-secreted proteins, cells were collected 2 days post-transfection and lysed in lysis buffer (50 mM Tris pH 7.6, 150 mM NaCl, 1 mM EDTA, 1% Triton-X). For secreted proteins, cell media was collected 5-7 days post-transfection. Protease inhibitor (Pierce, A32965) was added to all samples to avoid protein degradation. Westerns were conducted to confirm and roughly evaluate protein levels. Samples were stored at -80°C until use.

Anti-HA (EZView red Anti-HA affinity gel, Sigma 45-E6779), anti-V5 (Anti-V5 agarose affinity gel, Sigma 45-A7345), EZView red anti-c-Myc affinity gel (Sigma E6654), and EZView red Anti-FLAG beads (Sigma F2426) were used for immunoprecipitation (IP) of target proteins. In each trial, the two lysates (or media) each containing a protein of interest (POI) were mixed together before being added to appropriate beads. Single protein controls were applied directly to beads to assess for any non-specific binding. Samples were rotated on beads overnight at 4°C. The following day, unbound protein was removed. Five successive washes using wash buffer (50 mM Tris-HCl, pH 7.5, 150 mM NaCl, 1% Triton X-100) were done to remove any additional unbound proteins, with centrifugation to pellet beads between each wash. Bound proteins were eluted by the addition of elution buffer (100 mM Tris-HCl, pH 8.0, 1% SDS), followed by removal of beads using spin filters. Resulting eluate samples were prepared by the addition of 5x SDS buffer (0.2 M Tris⋅HCl, pH 6.8, 20% glycerol, 10% SDS, 0.25% bromophenol blue, 10% β-mercaptoethanol) and a 10 min incubation at 95°C.

### Sodium dodecyl sulfate-polyacrylamide gel electrophoresis (SDS-PAGE) and western blotting

Proteins from worm lysates or coIP experiments were separated on 10% or 4-20% gradient Mini-PROTEAN TGX Precast Gels (Bio-Rad Laboratories, Inc.) at 300V. Proteins were transferred to Immobilon-P PVDF membrane (MilliporeSigma) using Power Blotter Station (Model: PB0010, Invitrogen by Thermo Fisher Scientific) for 7 minutes for 10% gels or 8 minutes for 4-20% gels at 1.3A and 25V. Membranes were blocked with EveryBlot Blocking Buffer (Bio-Rad Laboratories, Inc.) for 5 minutes at room temperature. The resulting membranes were incubated with indicated primary antibody in Everyblot buffer at 4°C overnight with gentle shaking.

The following day, membranes were washed in 1xPBST (137 mM NaCl, 2.7 mM KCl, 10 mM Na_2_HPO4, 1.8 mM KH_2_PO4, 0.1% Tween-20) for 10 minutes, repeated three times total. Incubation with indicated secondary antibodies was done in Everyblot buffer or PBST + 5% dry milk (Carnation) for 2 hours at room temperature. After three additional PBST washes, membranes were developed using Clarity ECL Reagent (Bio-Rad 1705061) and imaged using a Bio-Rad ChemiDoc MP imaging system. When needed, membranes were stripped using Restore western blot stripping buffer (Pierce), blocked again, and reblotted.

Primary antibodies used include mouse monoclonal ANTI-FLAG® M2 antibody (diluted 1:5000, F3165, Sigma), mouse anti-HA IgG monoclonal antibody (Krackler, 12CAS, diluted 1:1000), rabbit anti-HA IgG monoclonal antibody (Cell Signaling, C29F4, diluted 1:1000), mouse anti-V5 IgG monoclonal antibody (Invitrogen (E10/V4RR), diluted 1:1000), rabbit anti-V5 IgG monoclonal antibody (Cell Signaling, D3H8Q, diluted 1:1000), mouse anti-actin IgM JLA20 monoclonal antibody (Developmental Studies Hybridoma Bank; diluted 1:2,000), and anti-Myc monoclonal antibody 9E10 (diluted 1:40). Secondary antibodies used include horseradish peroxidase-conjugated donkey anti-mouse IgG, peroxidase-conjugated goat anti-rabbit IgG, and horseradish peroxidase-conjugated goat anti-mouse IgM (all from Jackson ImmunoResearch; diluted 1:10,000).

### In silico protein-protein interaction structure predictions

The ColabFold [33] implementation of AlphaFold ([29] was used to predict structures of complexes involving different combinations of LON-2, SMOC-1, and the mature form of DBL-1. Default parameters were used except 3 models were predicted for each query. Pairwise combinations resulted in predictions for an interaction between LON-2 and SMOC-1, with an interface predicted template modeling (‘iptm’) score of 0.59 and an interaction between SMOC-1 and mature DBL-1 with an iptm score of 0.56. In contrast, no interaction was predicted between mature DBL-1 and LON-2. Given these predictions, we then attempted a prediction with LON-2, SMOC-1, and two copies of mature DBL-1 (mature DBL-1 is known to homodimerize). The result was a structural prediction in which SMOC-1 interacts with both mature DBL-1 and LON-2 (iptm score of 0.5) through interfaces that are the same as predicted in the two corresponding pairwise combinations. In each of these successful predictions, the three models predicted for each case had identical predicted interfaces. The iptm score is a confidence score generated by AlphaFold ([32]).

## ACKNOWLEDGEMENTS

We thank Claire Bénard, Andy Fire, Bob Goldstein, Erik Jorgensen, Oliver Hobert, and Daniel Zinshteyn for reagents, Herong Shi for amazing technical assistance before her untimely passing, Zhiyu Liu for generating the *jj244* allele, and the rest of the Liu lab for helpful discussions throughout the course of this work. We thank the Proteomics and Metabolomics Facility of Cornell University for providing the mass spectrometry data and NIH SIG grant 1S10 OD017992-01 support for the Orbitrap Fusion mass spectrometer. Some strains were obtained from the *C. elegans* Genetics Center, which is funded by NIH Office of 27 Research Infrastructure Programs (P40 OD010440). This work was supported by R35 GM136258 to JCF and R35 GM130351 to JL. MSD was partially supported by a National Science Foundation (NSF) Graduate Research Fellowship (DGE-1650441). MLMG and IL were students in the Molecular Biology and Genetics Research Experience for Undergraduate (MBG-REU) program, which was supported by NSF DBI1950247 and DBI1659534. TYC was partially supported by the Einhorn Discovery Grant and Undergraduate Research Fund in the College of Arts and Sciences at Cornell University.

