## Supplemental Figures and Tables for "*C. elegans* SMOC-1 interacts with both BMP and glypican to regulate BMP signaling"

**A**

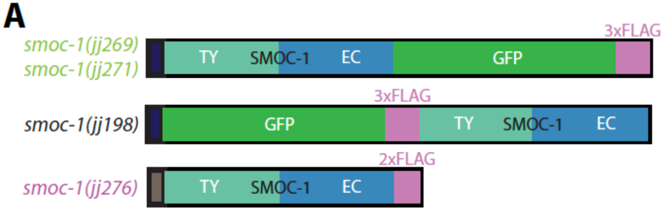


**B**

| Genotype | Susm penetrance (# of worms scored) |
|---|---|
| *sma-9(cc604)* | 1.5% (1359) [a] |
| *smoc-1(tm7125); sma-9(cc604)* | 98% (686) [b] |
| *smoc-1(jj198[GFP::3xFLAG::SMOC-1]); sma-9(cc604)* | 93% (328) |
| *smoc-1(jj269[SMOC-1::GFP::3xFLAG]); sma-9(cc604)* | 87% (1166) |
| *Smoc-1(jj276(SMOC-1::2xFLAG]); sma-9(cc604)* | 0% (723) |

**C**

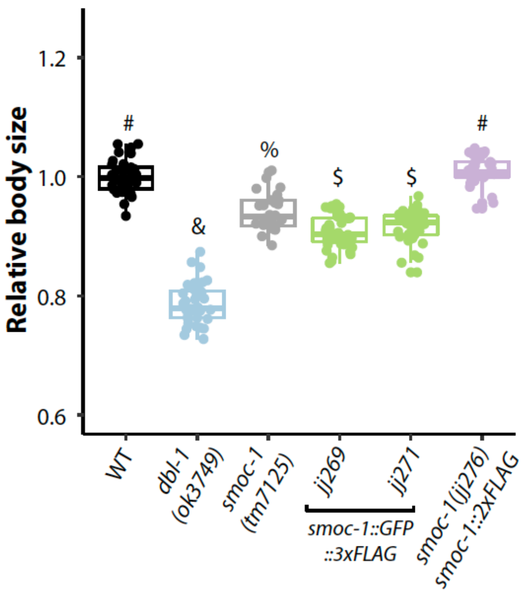


**Figure S1. GFP-tagged SMOC-1 is non-functional.**

**A)** Diagrams depicting various endogenously tagged SMOC-1 proteins, with the corresponding CRISPR alleles shown on the left of the diagrams. **B)** Table showing the penetrance of the Susm phenotype of strains carrying specified *smoc-1* allele in a *sma-9(cc604)* background. The Susm penetrance refers to the percent of animals with one or two M-derived CCs as scored using the *arIs37*(*secreted CC::GFP)* reporter. For each genotype, two independent isolates were generated (as shown in the strain list), four to seven plates of worms from each isolate were scored for the Susm phenotype at 20°C, and the Susm data from the two isolates were combined and presented in the table. [a] The lack of M-derived CCs phenotype is not fully penetrant in *sma-9(cc604)* mutants [1]. [b] Data from [2]. Statistical analysis was conducted by comparing various double mutants with the *sma-9(cc604)* single mutant. **** $P<0.0001$ (unpaired two-tailed Student's *t*-test). **C)** Relative body sizes of various strains at the same developmental stage (WT set to 1.0). Body sizes of 35 to 70 worms of each genotype were measured. A *dbl-1* null allele (*ok3749*) and *smoc-1* null allele (*tm7125*) were included as controls. Groups marked with distinct symbols are significantly different from each other ($P<0.001$, in all cases when there is a significant difference), while groups with the same symbol are not. Tested using an ANOVA with a Tukey HSD.

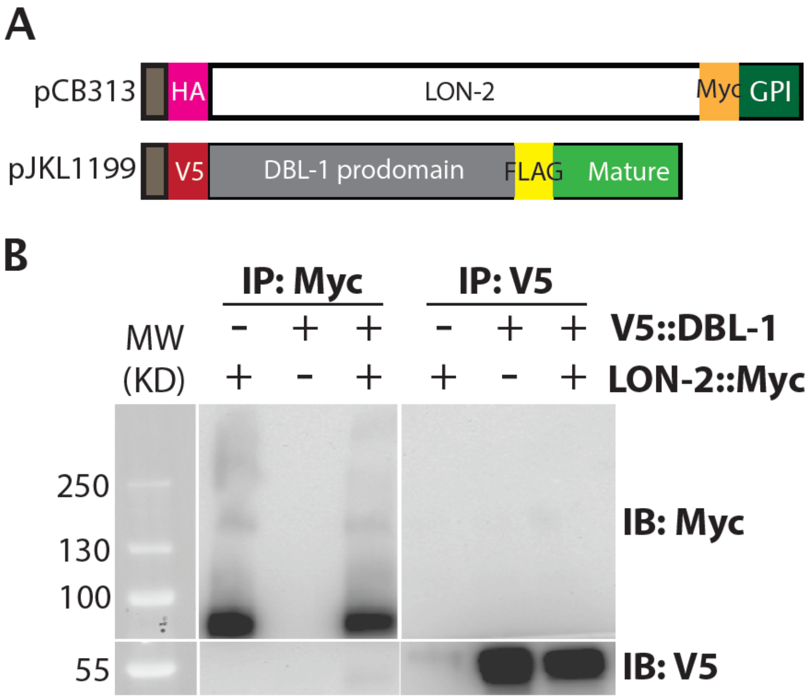


**Figure S2. LON-2 does not interact with either full length DBL-1.**
**A)** Diagrams of LON-2 and DBL-1 expression constructs used in the Drosophila S2 cell expression system. **B)** Results of co-IP experiments testing the interaction between HA::LON-2::Myc and V5::DBL-1 prodomain::FLAG::DBL-1 mature domain. Immunoprecipitation (IP) with anti-Myc beads or anti-V5 beads and immunoblot (IB) with anti-Myc or anti-V5 antibodies, as indicated. Full length DBL-1 detected by anti-V5 antibody runs at around 55KD. Experiments were independently repeated in triplicate, with representative results shown in this figure.

```
Ce-LON-2              -MVFR---------WLILFVLLYRSVLP----A--------------------------- 19
Cjp-LON-2             MHFFR---------WLILLVLLRSSTPT-EDAE--------------------------- 23
Cbn-LON-2             -MNFR---------WLILLVFYFRSISP----T--------------------------- 19
Cbr-LON-2             -MNFR---------WLILLVFLFRTALP---LS--------------------------- 20
Cre-LON-2             -MNFR---------WLILLVLLFRTALS----S--------------------------- 19
Drosophila-Dally      -MAARSVRLAQLLLFTLLCGFVGLSAAKHLDLDGIHHHQHHLHSATTHHRRRLQRDSRAK 59
Mouse-Glypican        -MELR------TRGWWLLCAAAALVVCA-------------------------------R 22
Human-Glypican        -MELR------ARGWWLLCAAAALVACA-------------------------------R 22
                          *         : :*

Ce-LON-2              ----------EEV-VVV---DIL-----TNS---TSLEEPTEEYTC-DCNTDD-LIQKGN 55
Cjp-LON-2             ----------NEK-VVI---DLL-----TNT---TTVEE-SEDVIC-DCNGSD-MLEKAN 58
Cbn-LON-2             ----------DDA-VVF---DIL-----TNS---TSLEEPTEEYTC-ECDVEE-LTEKGN 55
Cbr-LON-2             ----------EEV-VIV---DTL-----TNS---TALEEPTDEYTC-DCNDKD-LLEKGN 56
Cre-LON-2             ----------EEV-VVV---DIL-----TNS---TSLEEPTDEFTC-ECDAED-LLEKGN 55
Drosophila-Dally      DAVGGSTHQCDAVKSYFESIDIKSSG--TYSEKGRHLRRKL--LQQCHGAGA---AGQSR 112
Mouse-Glypican        GDPASKSRSCSEVRQIYGAKGFSLSDVPQAEISGEHLRICPQGYTCCTSEMEENLANHSR 82
Human-Glypican        GDPASKSRSCGEVRQIYGAKGFSLSDVPQAEISGEHLRICPQGYTCCTSEMEENLANRSH 82
                                          .               :.                   :..

                                                                   $$
Ce-LON-2              YTLTVEVKNMQVRLVEPFRDAIAFTTGEKKHLLDFIRFVTLREVRSTYPLLLNYTDFLNS 115
Cjp-LON-2             YTLTVEVKNMQVRLVEPFRDAIAFTTSEKQHLLRFVKYVTLRDIRSTYPMLLNYTDFYAN 118
Cbn-LON-2             YTLTIEVRNMQVRLVEPFRDAIAFVASEKQHLLDFIKFVTLRDIKADYPLLLNYTEFYDS 115
Cbr-LON-2             YTLTVEVKNMQVRLVEPFRDAIAFITSEKQHLLDYIQFVVLRDTKATYPLLLNYTEYASS 116
Cre-LON-2             YTLTVELRNMQVRLVEPFRDAIAFVASEKTHLLDFIKFVTLRDIRSTYPLLLNYTEFYTN  115
Drosophila-Dally      RNVRATA--SSP-HQQPARSPRDECQSHVLELAQISE----NMTHSLFSK--VYTRMVPS 163
Mouse-Glypican        MELESALHDSSRALQATLATQLHGIDDHFQRLLNDSE----RTLQEAFPG--AFGDLYTQ 136
Human-Glypican        AELETALRDSSRVLQAMLATQLRSFDDHFQHLLNDSE----RTLQATFPG--AFGELYTQ 136
                        :       .               ..  .*    .    .  :  :     :     .

                                                                      $  $
Ce-LON-2              ----FDEMIGTFRNILT-------TE---NAISLTGIKYEVSTAVQKFLSAILPDMFLCL 161
Cjp-LON-2             ----YDELLGTIANILL-------TE---NDISFSGIRYEVTTAVDKFLKSLLPDMFLCL 164
Cbn-LON-2             ----FDGLIGSLRGILT-------TD---NAISLSGIKYEVSTAVQKFLNKLLPDMFLCL 161
Cbr-LON-2             ----FDELIKTFKNILS-------TE---NDILLSGIKYEVTTAVQKFFTSLLPDMFLCL 162
Cre-LON-2             ----FDDLIGTFRNILT-------TE---NAISLSGIKYEVTTAVQKFLSALLPDMFLCL 161
Drosophila-Dally      SRMMIHQLYTEIMNHLIYTSNYTNSNGQLGRRGIGSVQSNLEEAVRHFFVQLFPVAYHQM 223
Mouse-Glypican        NTRAFRDLYAELR---------------LY---YRGANLHLEETLAEFWARLLERLFKQL 178
Human-Glypican        NARAFRDLYSELR---------------LY---YRGANLHLEETLAEFWARLLERLFKQL 178
                             :   :                       . . .:  ::  .*   ::   :  :

                      $     $       $
Ce-LON-2              S---VGKCRTVPLDYHNCMMASTKHWSVYLGNTPNKMAMTISEAIYRYRKVEFLLV---D 215
Cjp-LON-2             S---VGKCRTVPLEYHNCMLENTKHWSTYIGNTPNKMAATISDTIYRYRKVEFLLV---D 218
Cbn-LON-2             S---VGKCRTVPSDYYNCMMASTEHWSVYLGNTPNKMAMTVSEAIYRYRKVEFLLV---D 215
Cbr-LON-2             S---VGKCRTVPLEYQNCMTASTEHWSVYIGNTPNKMAVTIAEAIYRYRKVEFLLV---D 216
Cre-LON-2             S---VGKCRTVPLEYHNCMMTSTEHWSEYIGNTPNKMAMTIADAIYRYRKVEFVLV---D 215
Drosophila-Dally      VHLSKNNLGDLHEDYVNCLQHNFDEMHP-FGDIPQQVQSNLGKSVHMSNVFMNALLQAAE 282
Mouse-Glypican        H-----PQL-LPDDYLDCLGKQAEALRP-FGDAPRELRLRATRAFVAARSFVQGLGVASD 231
Human-Glypican        H-----PQLLLPDDYLDCLGKQAEALRP-FGEAPRELRLRATRAFVAARSFVQGLGVASD 232
                                :  :* :*:  . .     :*: *.::      :.   . .   *    :

Ce-LON-2              MHKQLMNAHNLTLSHECLQEYVHTLPCN-C------TMAGITPCHTSCSNSMEKCFGKFS 268
Cjp-LON-2             MHKQLMNAHNLTLTDACLQEYVTTLPCN-C------TTSGIGRCKTSCSDAMEKCFGKYS 271
Cbn-LON-2             MHRQLMQAHSLTITDECLSEYVNTLPCN-C------TMLGIARCGTSCPNSMEKCFGKFS 268
Cbr-LON-2             MHQQLMNAQSLTITDECLKQYVHSLPCN-C------TLSGIVRCQTSCATSMETCFGKYS 269
Cre-LON-2             MHKQLMNAHSLTVTDGCLTEYVTTLPCN-C------TMLGIVRCHTSCSDSMETCFGKYS 268
Drosophila-Dally      VLSEADALYGEQLTDTCKLHLLKMHYCPNCNGHHSSSRSETKLCYGYCKNVMRGCSAEYA 342
Mouse-Glypican        VVRK---VAQVPLAPECSRAIMKLVYCAHCRGVPG-----ARPCPDYCRNVLKGCLANQA 283
Human-Glypican        VVRK---VAQVPLGPECSRAVMKLVYCAHCLGVPG-----ARPCPDYCRNVLKGCLANQA 284
                      :  :        :   *    :    *  *             *   *   :. * .: :
```

```
                                                                   $$  $$      $
Ce-LON-2              ----REWAAKLHLMRNMTSTKKSFLDEFLSLK---KTIFSVIRIFIERKSYVYAEHVFNS 321
Cjp-LON-2             ----RDWSAKLHLMRNMTNTKKRFLEGFLTLK---KTIFSAIRLFIERKSFVYAEHVFNS 324
Cbn-LON-2             ----REWAAKLHLMRNMTNSKKSYLEEFLTLK---KTIFQVIRQFIERKSFVYAEHVFNS 321
Cbr-LON-2             ----REWAAKLHLMRNMTSTKKSFLDEFLTLR---KTIFSVIRLFIERKSFEYAAYVYKA 322
Cre-LON-2             ----REWAAKLHLMKNMTSTKKSFLEEFLTLK---KTIFSVIRLFIERKSFLYAEQVFKS 321
Drosophila-Dally      GLLDSPWSGVVDSLNNLVTTHILSDTGIINVIKHLQTYFSEAIMAAMHNGPELEKKVKKT 402
Mouse-Glypican        D-LDAEWRNLLDSMV-LITDKFWGPSGAESVIGGVHVWLAEAINALQDNKDTLTAKVIQA 341
Human-Glypican        D-LDAEWRNLLDSMV-LITDKFWGTSGVESVIGSVHTWLAEAINALQDNRDTLTAKVIQG 342
                            *   :. :  : . :        .:    :. :         :       * :

                         $
Ce-LON-2              CGPLGEMIIH-------PSKHS-----VHFQSPGPFVSRGDGAVRELQLSAKTWDRLGRK 369
Cjp-LON-2             CGPLGEMIIH-------PSKHI-----FHLFSPGPFVSRADGAVRELQLSTKMWEKLGRK 372
Cbn-LON-2             CGPLGQMIIH-------PSKHV-----TRFHSPGPFVSRADGAVKELQLSAKSWDRFGRK 369
Cbr-LON-2             CGPLGEMIIH-------PSKHV-----ARIHSPGPFVSRADGAVRELQLSAKSWDRFGRK 370
Cre-LON-2             CGPLGEMIIH-------PSKHV-----AQIHSPGPFVSRADNAVRELQLSAKSWDRFGRK 369
Drosophila-Dally      CGTPSLTPYSSGEPDARPPPHKNNVKWATDPDPGMVLF-----LSTIDKSKEFYTTIVDN 457
Mouse-Glypican        CGNPKVNPHGSGPEEKRRRG---KLALQEKPSTGTLEKLVSEAKAQLRDIQDFWISLPGT 398
Human-Glypican        CGNPKVNPQGPGPEEKRRRG---KLAPRERPPSGTLEKLVSEAKAQLRDVQDFWISLPGT 399
                      **                               * .          :    . :  :  .

Ce-LON-2              ICDHSGV--VLNPTMCYDGTKVISIDHDLL-PITKDVRPRPMMDWIEKK----------- 415
Cjp-LON-2             ICEHNGI--VYHPTLCFDGTKVISIEHELP-PITKDVRPRPMMDWIEKR----------- 418
Cbn-LON-2             ICDH-GV--VLHPGMCYDGTKVIT------------------------------------ 390
Cbr-LON-2             ICDHTGV--VVHPGMCYDGTKVIKVEHELL-PITKDVRPRPMMDWIEKK----------- 416
Cre-LON-2             ICDHTGV--VLHPGYCFDGTKVIKIQHELA-PITKDVRPKTMMDWIEKK----------- 415
Drosophila-Dally      FCDEQQH--SRDDHSCWSGDRFGDYTQLLINPGTDSQRYNPEVPFNAKAQTGKLNELVDK 515
Mouse-Glypican        LCSEKMAMSPASDDRCWNGISKGRYLPEVMGDGLANQINNPEVEVDITKPDMTIRQQIMQ 458
Human-Glypican        LCSEKMALSTASDDRCWNGMARGRYLPEVMGDGLANQINNPEVEVDITKPDMTIRQQIMQ 459
                      :*..           *:.*

                                                     &&&&
Ce-LON-2              ---NEKKA--NVEGSASNPLWDDEDSEDFDGSGSGMPPV-IDRNPVKAIIQEDHPKNIDL 469
Cjp-LON-2             ---GEKKPTAVVEGS-ASPLWDDEDSDDFEGSGSGLPTTIDNKEPTKT---EEPTKTIDL 471
Cbn-LON-2             ------------------------------------------------------------ 390
Cbr-LON-2             ---NEKKI--AVEGSA-SPLWDDEDSDGFEGSGSGMPPIIIDRNPVKAVIQEDHPKNIDL 470
Cre-LON-2             ---NEKKI--AVEGSA-SPLWDDEDFDGFEGSGSGMPPNIIDRNPVKAVIQQDHPKNIDL 469
Drosophila-Dally      LFKIRKSIG-AAAPSNSIQATHDIQNDMGEGSGGGEGQIGDDEEEYGGAHGSG-DGSGDG 573
Mouse-Glypican        LKIMTNRLR-GAYGGNDVDFQD--ASDDGSGSGSGGGCPDDTC---GRRVSKK-SS---S 508
Human-Glypican        LKIMTNRLR-SAYNGNDVDFQD--ASDDGSGSGSGDGCLDDLC---GRKVSRK-SS---S 509

Ce-LON-2              STNPKGPS------VIVTEKEIQPDGSTTSSILICVII-VAVIKLF--------       508
Cjp-LON-2             SINPKGPS------VIANEQENSPEPDGSSTICPITIL-ALVLAFWCI------       512
Cbn-LON-2             ------------------------------------------------------       390
Cbr-LON-2             STNPKGPS------VIVTEQGSQPDGSSTVSALTSLII-IAIIRFF--------       509
Cre-LON-2             STNPKGPS------VIVTEQGVQPDGSSILTIFTSLIF-IAIIRLF--------       508
Drosophila-Dally      PHTPIEESEGTTTNEVESRDSGKTSGSNPLEGTATWMLLTLVTMLFSSCS----       623
Mouse-Glypican        SRTPLTHALP----GL-SEQEGQKTSAATCPEPHSFFLLFLVTLVLAAARPRWR       557
Human-Glypican        SRTPLTHALP----GL-SEQEGQKTSAASCPQPPTFLLPLLLFLALTVARPRWR       558
```

**Figure S3. Multiple sequence alignment of LON-2 homologs.**
Clustal Omega (CLUSTAL O(1.2.4)) [3] of *C. elegans* (Ce) LON-2 with its homologs from other nematode species, including *C. Japonica* (Cjp), *C. brenneri* (Cbn), *C. briggsae* (Cbr) and *C. remanei* (Cre), as well as with Drosophila Dally and Glypican from *M. musculus* (mouse) and *H. sapiens* (human). $ marks the residues at the interface between LON-2::SMOC-1, as identified via ColabFold [4]. &&&& marks the glycosaminoglycan attachment site.

```
                                  $    $ $$$$$ $
C.elegans_DBL-1         CRRTDFYVDFDDLNWQDWIMAPKGYDAYQCQGSCPNPMPAQLNATNHAIIQSLLHSLRPD  60
C.brenneri_DBL-1        CRRTDLYVDFDDLGWQDWIMAPKGYDAYQCQGSCPNPMPAQLNATNHAIIQSLLHSIKPD  60
C.remanei_DBL-1         CRRTDLYVDFDDLGWQDWIMAPKGYDAYQCQGSCPNPMPAHLNATNHAIIQSLLHSLKPD  60
C.briggsae_DBL-1        CRRTDLYVDFDDLGWQDWIMAPKGYDAYQCQGSCPNPMPAQLNATNHAIIQSLLHSLKPD  60
D.melanogaster_DPP      CRRHSLYVDFSDVGWDDWIVAPLGYDAYYCHGKCPFPLADHFNSTNHAVVQTLVNNMNPG  60
H.sapiens_BMP2          VGDPPLLVNFSDVGWNDWIVAPPGYHAFYCHGECPFPLADHLNSTNHAIVQTLVNSVN-S  59
X.laevis_BMP4           CRRHSLYVDFSDVGWNDWIVAPPGYQAFYCHGDCPFPLADHLNSTNHAIVQTLVNSVN-S  59
H.sapiens_BMP4          CRRHSLYVDFSDVGWNDWIVAPPGYQAFYCHGDCPFPLADHLNSTNHAIVQTLVNSVN-S  59
X.laevis_BMP2           CRRHPLYVDFSDVGWNDWIVAPPGYHAFYCHGECPFPLADHLNSTNHAIVQTLVNNVN-P  59
                             : *:*.*:.*:***:** **.*: *:*.** *:  ::*:****::*:*::.:.

                                        $$$$    $$$$$$$$ $
C.elegans_DBL-1         EVPPPCCVPTETSPLSILYMDVDKVIVIREYADMRVESCGCR  102
C.brenneri_DBL-1        EVPPPCCVPTETSPLSILYMDVDKVIVIREYADMRVDSCGCR  102
C.remanei_DBL-1         EVPPPCCVPTETSPLSILYMDVDKVIVIREYADMRVDSCGCR  102
C.briggsae_DBL-1        EVPPPCCVPTETSPLSILYMDVDKVIVIREYADMRVDSCGCR  102
D.melanogaster_DPP      KVPKACCVPTQLDSVAMLYLNDQSTVVLKNYQEMTVVGCGCR  102
H.sapiens_BMP2          KIPKACCVPTELSAISMLYLDENEKVVLKNYQDMVVEGCGCR  101
X.laevis_BMP4           SIPKACCVPTELSAISMLYLDEYDKVVLKNYQEMVVEGCGCR  101
H.sapiens_BMP4          SIPKACCVPTELSAISMLYLDEYDKVVLKNYQEMVVEGCGCR  101
X.laevis_BMP2           NIPKACCVPTELSAISMLYLDENEKVVLKNYQDMVVEGCGCR  101
                        .:*  *****: . :::**::  . :*:::* :* * .****
```

**Figure S4. Multiple sequence alignment of the mature domains of DBL-1 homologs.** Clustal Omega (CLUSTAL O(1.2.4)) alignment [3] of the mature domains of DBL-1 with its homologs from other nematode species, as well as homologs from Drosophila, Xenopus and humans. $ marks the residues at the interface between mature DBL-1 and SMOC-1, as identified by ColabFold [4]**.** The first cysteine residue in the mature domain of DBL-1 homologs is assigned as the #1 position.

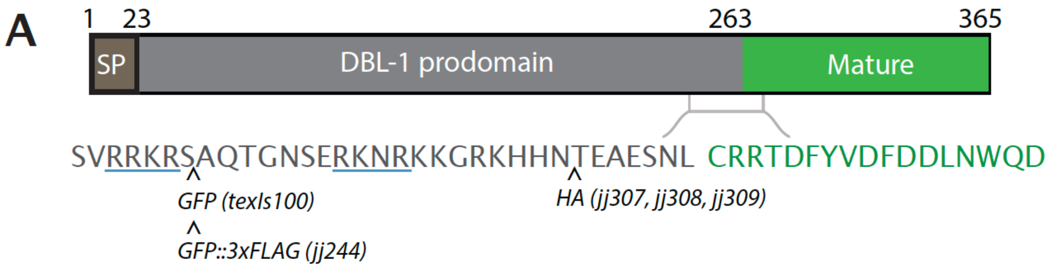


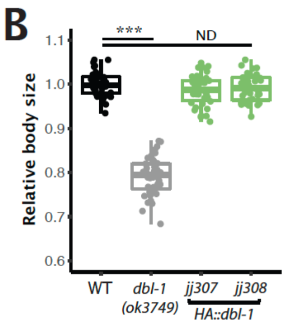


C

| Genotype | Susm penetrance (# of worms scored) |
|---|---|
| *sma-9(cc604)* | 1.5% (1359)[a] |
| *dbl-1(nk3); sma-9(cc604)* | 100% (55)[b] *** |
| *dbl-1(wk70); sma-9(cc604)* | 100% (60)[b] *** |
| *dbl-1(jj308); sma-9(cc604)* | 3.1% (387)[ND] |

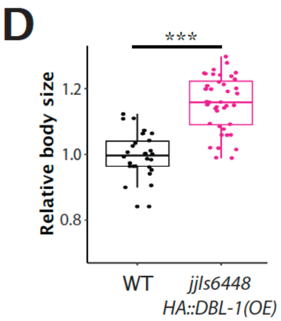


**Figure S5: Endogenously tagged HA::DBL-1 is fully functional, and when over-expressed, HA::DBL-1 can cause a long body size phenotype.**

**A)** Schematic of DBL-1 protein, which contains a signal peptide (SP, dark grey), a prodomain (light grey), and a cysteine-knot containing mature domain that is the active signaling ligand (green). Inset shows residues flanking the prodomain-mature domain boundary, indicating the predicted furin cleavage sites (underlined residues) in relation to the placements of the tags (arrows). *texIs100* is an integrated transgene over-expressing a GFP-tagged DBL-1 [5]. However, when GFP was inserted in the same location as in *texIs100* in the endogenous *dbl-1* locus, the resulting allele, *jj244*, is not functional. *jj307, jj308* and *jj309* are three identical alleles generated via CRISPR, with the HA tag inserted in the marked location. **B)** Relative body lengths of synchronized larvae stage-matched at L4.3 vulva stage grown at 20°C, with the body length of WT worms set to 1.0. Sample sizes are 40 for each genotype. An ANOVA followed by Tukey's honest significant difference was used to test for differences between genotypes. ****P*<0.001; ND, no difference. **C)** Table showing the penetrance of the Susm phenotype of double mutant strains between *sma-9(cc604)* and different *dbl-1* alleles. [a] The lack of M-derived CCs phenotype is not fully penetrant in *sma-9(cc604)* mutants. [b] Data for the two null *dbl-1* alleles, *nk3* and *wk7*, are from [1]. For *jj308*, two independent isolates were generated, and the Susm data from the two isolates were combined and presented in the table. Statistical analysis was conducted by comparing the strains carrying *dbl-1* alleles with the *sma-9(cc604)* single mutants. *** *P*<0.001 (unpaired two-tailed Student's t-test). ND, no difference. **D)** Relative body lengths of stage-matched WT worms (set to 1.0) and worms carrying an integrated transgene (*jjIs6448*) that overexpresses HA::DBL-1. WT: N=35. *jjIs6446*, N=41. ****P*<0.001 (ANOVA followed by Tukey HSD). **E)** Schematic of GFP::3xFLAG::LON-2 protein in *lon-2(jj207)* animals.

**Table S1: LON-2 peptides recovered from the IP-MS experiments.** 1) IP using full

length SMOC-1::2xFLAG (experiment 1)

| |
|---|
| [K].MAMTISEAIYR.[Y] |
| [R].NILTTENAISLTGIK.[Y] |
| [K].VISIDHDLLPITK.[D] |
| [K].SFLDEFLSLK.[K] |
| [R].KICDHSGVVLNPTMCYDGTK.[V] |
| [K].HWSVYLGNTPNK.[M] |
| [K].HSVHFQSPGPFVSR.[G] |
| [K].ICDHSGVVLNPTMCYDGTK.[V] |
| [K].ICDHSGVVLNPTMCYDGTK.[V] |
| [R].KVEFLLVDMHK.[Q] |
| [R].DAIAFTTGEK.[K] |
| [R].NILTTENAISLTGIK.[Y] |
| [K].GPSVIVTEK.[E] |

2) IP using full length SMOC-1::2xFLAG (experiment 2)

| |
|---|
| [R].DAIAFTTGEK.[K] |
| [K].GPSVIVTEK.[E] |
| [K].HSVHFQSPGPFVSR.[G] |
| [K].HWSVYLGNTPNK.[M] |
| [R].KICDHSGVVLNPTMCYDGTK.[V] |
| [R].KVEFLLVDMHK.[Q] |
| [K].ICDHSGVVLNPTMCYDGTK.[V] |
| [K].ICDHSGVVLNPTMCYDGTK.[V] |
| [K].MAMTISEAIYR.[Y] |
| [K].NIDLSTNPK.[G] |
| [R].NILTTENAISLTGIK.[Y] |
| [R].NILTTENAISLTGIK.[Y] |
| [K].SFLDEFLSLK.[K] |
| [K].SYVYAEHVFNSCGPLGEMIIHPSK.[H] |
| [K].SYVYAEHVFNSCGPLGEMIIHPSK.[H] |
| [K].SYVYAEHVFNSCGPLGEMIIHPSK.[H] |
| [K].VISIDHDLLPITK.[D] |
| [K].YEVSTAVQK.[F] |

**3) IP using SMOC-1(EC)::2xFLAG**

| |
|---|
| [R].DAIAFTTGEK.[K] |
| [K].GPSVIVTEK.[E] |
| [K].HSVHFQSPGPFVSR.[G] |
| [K].HWSVYLGNTPNK.[M] |
| [R].KICDHSGVVLNPTMCYDGTK.[V] |
| [R].KVEFLLVDMHK.[Q] |
| [K].ICDHSGVVLNPTMCYDGTK.[V] |
| [K].ICDHSGVVLNPTMCYDGTK.[V] |
| [K].MAMTISEAIYR.[Y] |
| [K].NIDLSTNPK.[G] |
| [R].NILTTENAISLTGIK.[Y] |
| [K].SFLDEFLSLK.[K] |
| [K].SYVYAEHVFNSCGPLGEMIIHPSK.[H] |
| [K].VISIDHDLLPITK.[D] |
| [K].YEVSTAVQK.[F] |

**Table S2. Plasmids generated in this study.**

| Plasmid ID | Details |
|---|---|
| **Plasmids for expression in *C. elegans*** | |
| pMSD35 | *smoc-1p::smoc-1 gDNA::2xflag::smoc-1 3'UTR* |
| pMSD43 | *smoc-1p::smoc-1(D219N, D223N) gDNA::2xflag::smoc-1 3'UTR* |
| pMSD44 | *smoc-1p::smoc-1 (TY) gDNA::2xflag::smoc-1 3'UTR* |
| pMSD45 | *smoc-1p::smoc-1(EC) gDNA::2xflag::smoc-1 3'UTR* |
| pMSD46 | *smoc-1p::smoc-1(jj65(C210Y)) gDNA::2xflag::smoc-1 3'UTR* |
| pMSD47 | *smoc-1p::smoc-1(jj85(E105K)) gDNA::2xflag::smoc-1 3'UTR* |
| pMSD58 | *smoc-1p::smoc-1 gDNA:: 2xflag::TM::smoc-1 3'UTR* |
| pMSD60 | *smoc-1p::smoc-1(D219N, D223N, D229, E230N) gDNA::2xflag::smoc-1 3'UTR* |
| pMSD64 | *smoc-1p::hsmoc1(EC) cDNA::2xflag::smoc-1 3'UTR* |
| pMSD65 | *smoc-1p::hsmoc2(EC) cDNA::2xflag::smoc-1 3'UTR* |
| pMSD71 | *smoc-1p::smoc-1 (EC) gDNA:: 2xflag::TM::smoc-1 3'UTR* |
| pMSD1 | sgRNA plasmid #1 for generating *smoc-1::2xflag* and *smoc-1(EC)::2xflag (in pRB1017)*<br>MSD-1: TCTTGATTCTGATCTTAAAATGTAC<br>MSD-2: AAACGTACATTTTAAGATCAGAATC |
| pMSD2 | sgRNA plasmid #2 for *smoc-1::2xflag* and *smoc-1(EC)::2xflag (in pRB1017)*<br>MSD-3: TCTTGAACATTGCAAATTGAGGGGG<br>MSD-4: AAACCCCCCTCAATTTGCAATGTTC |
| pMSD67 | sgRNA plasmid #1 for generating *smoc-1(TY)::2xflag (in pRB1017)*<br>MSD-163: TCTTGAGAAGAGAACACGATTTCTG<br>MSD-164: AAACCAGAAATCGTGTTCTCTTCTC |
| pMSD68 | sgRNA plasmid #2 for generating *smoc-1(TY)::2xflag (in pRB1017)*<br>MSD-165: TCTTGGAGAAGAAACAATCGGTGTA<br>MSD-166: AAACTACACCGATTGTTTCTTCTCC |
| pMSD78 | sgRNA plasmid #3 for generating *smoc-1(TY)::2xflag (in pRB1017)*<br>MSD-187: TCTTGGAAGGAGCATCGGGATCCAG<br>MSD-188: AAACCTGGATCCCGATGCTCCTTCC |
| pMSD79 | sgRNA plasmid #4 for generating *smoc-1(TY)::2xflag (in pRB1017)*<br>MSD-189: TCTTGCTGTACATTTTAAGATCAGA<br>MSD-190: AAACTCTGATCTTAAAATGTACAGC |
| pMSD84 | Repair template for generating *smoc-1(EC)::2xflag* |
| pJKL1204 | sgRNA plasmid #1 for generating *HA::dbl-1 (in pRB1017)*<br>JKL-1840: TCTTGGTAGAAAGCATCATAACACCG<br>JKL-1841: AAACCGGTGTTATGATGCTTTCTAC |
| pJKL1205 | sgRNA plasmid #2 for generating *HA::dbl-1 (in pRB1017)*<br>JKL-1842: TCTTGCCTCCGACAAAGATTGCTCT<br>JKL-1843: AAACAGAGCAATCTTTGTCGGAGG |
| pTYC3 | *dbl-1p::HA::dbl-1 gDNA::dbl-1 3'UTR* |

| **Plasmids for expression in *Drosophila* S2 cells** | |
|---|---|
| pJKL1210 | *pAc5::sax-7ss::smoc-1::V5* |
| pJKL1211 | *pAc5::sax-7ss::HA::smoc-1* |
| pMSD49 | *pAc5::sax-7ss::smoc-1(TY)::V5* |
| pMSD50 | *pAc5::sax-7ss::smoc-1(EC)::V5* |
| pMSD54 | *pAc5::sax-7ss::HA::smoc-1(TY)* |
| pMSD55 | *pAc5::sax-7ss::HA::smoc-1(EC)* |
| pCB313 | *pAc5::sax-7ss::HA::lon-2::myc (gift from C. Benard)* |
| pJKL1241 | *pAc5::sax-7ss::lon-2::myc* |
| pJKL1198 | *pAc5::HA::dbl-1pro::FLAG::dbl-1mature* |
| pJKL1199 | *pAc5::V5::dbl-1pro::FLAG::dbl-1mature* |

All plasmids were verified by Sanger sequencing.

**Table S3. Oligonucleotides used in this study**

| Oligo ID | Sequence |
|---|---|
| ***smoc-1::2xflag repair oligo*** | |
| MSD-67 | CCAGCCAAACGTCCAGATCAACTAAACCCATTTCTGTACATTTTAAGATCAGAAGGAGCATCGGGATCCAGTGGA GCATCGGATTATAAAGACGATGACGATAAGCGTGACTACAAGGACGACGACGACAAGCGTTAATTTTAAGTTTTA ATTCTCCCCCTCAATTTGCAATGTTCTTTAAAAATCTACCA |
| ***smoc-1(TY)::2xflag repair oligo*** | |
| MSD-191 | GTGAAGAAACTCCAACAACTACAGTTGCTCCAAAAAGAGTGAGAAGAGGAGCTTCCGGTTCTAGTGGAGCATCG GATTATAAAGACGATGACG |
| ***HA::dbl-1 repair oligo*** | |
| MSD-110 | TTGTAGGGTAGAAAGCATCATAACACCGAAGCTGAGGGATCCAGTGGAGCATCGTACCCATACGACGTCCCAGA CTACGCCGGAGCATCGGGATCCAGTAGCAATCTTTGTCGGAGGACTGATTTCTACGTGG |
| **For amplification of *smoc-1::2xflag* from *jj276*** | |
| JKL-1549 | ATCTAGCCCGGGTTTCCCCCATCTACAATCATCCAAGTTTTG |
| JKL-1550 | TATCTCGGGCCCTTATTGCGAATGATAAACCCATTCAAGCTG |
| **For genotyping *jj276*** | |
| MSD-10 | AGAATGTCAGACAGTGCTCC |
| MSD-70 | GACTACAAGGACGACGACGA |
| JKL-1211 | TAATGGAAGGAGGTTACCCG |
| **For genotyping *jj411/412*** | |
| MSD-10 | AGAATGTCAGACAGTGCTCC |
| MSD-28 | GCCAAAGTTAGGCTCATCGACAACAAGAGG |
| JKL-1211 | TAATGGAAGGAGGTTACCCG |
| **For genotyping *jj441*** | |
| JKL-1517 | AGAAAGGGGGACTTGGTAGG |
| JKL-1519 | GGTTTCCATTCACTTCTTTCAAGCAC |
| MSD-66 | CGTATACATATTTGTTAAGTTTACTCAATTTTCAG |
| **For genotyping *ok4128*** | |
| JKL-1981 | CTGACAAGCCAGTTCACGAG |
| JKL-1982 | CCCAGATTTCGGAACCTCAC |
| JKL-1983 | CAACAAACCACCAACACATGAG |
| **For genotyping *jj307/8/9*** | |
| ZL-499 | AGACACAGAGCAGCTCACTGAGC |
| MSD-99 | GATAAGCATCGTAGCCCTCTG |

**Table S4.** ***C. elegans*** **strains used in this study**

| Strain ID | Genotype |
|---|---|
| **Strains carrying endogenously tagged SMOC-1::2xFLAG** | |
| LW5524 | *smoc-1(jj276[smoc-1::2xflag]) V* |
| LW5525 | *arIs37[secreted CC::gfp] I; smoc-1(jj276[smoc-1::2xflag]) V; sma-9(cc604) X, isolate #1* |
| LW5527 | *arIs37[secreted CC::gfp] I; smoc-1(jj276[smoc-1::2xflag]) V; sma-9(cc604) X, isolate #2* |
| **Strains overexpressing untagged SMOC-1** | |
| LW5130 | *jjIs5119[pMSD4.4(smoc-1p::smoc-1) + LiuFD188(myo-2p::mCherry)], x3* |
| **Strains overexpressing SMOC-1::2xFLAG** | |
| LW5798 | *jjIs5798[pMSD35.7(smoc-1p::smoc-1::2xflag) + LiuFD290(ttx-3p::RFP)], x0* |
| LW5799 | *jjIs5799[pMSD35.7(smoc-1p::smoc-1::2xflag) + LiuFD290(ttx-3p::RFP)], x0* |
| LW5800 | *jjIs5800[pMSD35.7(smoc-1p::smoc-1::2xflag) + LiuFD290(ttx-3p::RFP)], x0* |
| LW5812 | *jjIs5798[pMSD35.7(smoc-1p::smoc-1::2xflag) + LiuFD290(ttx-3p::RFP)], x2 isolate #1* |
| LW5813 | *jjIs5798[pMSD35.7(smoc-1p::smoc-1::2xflag) + LiuFD290(ttx-3p::RFP)], x2 isolate #2* |
| LW5814 | *jjIs5799[pMSD35.7(smoc-1p::smoc-1::2xflag) + LiuFD290(ttx-3p::RFP)], x2 isolate #1* |
| LW5815 | *jjIs5799[pMSD35.7(smoc-1p::smoc-1::2xflag) + LiuFD290(ttx-3p::RFP)], x2 isolate #2* |
| LW5816 | *jjIs5800[pMSD35.7(smoc-1p::smoc-1::2xflag) + LiuFD290(ttx-3p::RFP)], x2 isolate #1* |
| LW5817 | *jjIs5800[pMSD35.7(smoc-1p::smoc-1::2xflag) + LiuFD290(ttx-3p::RFP)], x2 isolate #2* |
| LW6061 | *jjEx6061[pMSD35.7(smoc-1::2xflag) + LiuFD290(ttx-3p::RFP)]* |
| LW6062 | *jjEx6062[pMSD35.7(smoc-1::2xflag) + LiuFD290(ttx-3p::RFP)]* |
| LW6087 | *jjEx6087[pMSD35.7(smoc-1::2xflag) + LiuFD290(ttx-3p::RFP)]; smoc-1(tm7125) V* |
| LW6088 | *jjEx6088[pMSD35.7(smoc-1::2xflag) + LiuFD290(ttx-3p::RFP)]; smoc-1(tm7125) V* |
| LW6091 | *jjEx6091[pMSD35.7(smoc-1::2xflag) + LiuFD290(ttx-3p::RFP)]; arIs37[secreted CC::gfp] I; cup-5(ar465) III; smoc-1(tm7125) V; sma-9(cc604) X* |
| LW6109 | *jjEx6109[pMSD35.7(smoc-1::2xflag) + LiuFD290(ttx-3p::RFP)]; arIs37[secreted CC::gfp] I; cup-5(ar465) III; smoc-1(tm7125) V; sma-9(cc604) X* |
| LW6110 | *jjEx6110[pMSD35.7(smoc-1::2xflag) + LiuFD290(ttx-3p::RFP)]; arIs37[secreted CC::gfp] I; cup-5(ar465) III; smoc-1(tm7125) V; sma-9(cc604) X* |
| **Strains carrying endogenously tagged SMOC-1 truncations** | |
| LW6276 | *smoc-1(jj411[smoc-1(TY)::2xflag]) V* |
| LW6277 | *smoc-1(jj412[smoc-1(TY)::2xflag]) V* |
| LW6427 | *arIs37[secreted CC::gfp] I; cup-5(ar465) III; smoc-1(jj411[smoc-1(TY)::2xflag]) V; sma-9(cc604) X isolate #1* |
| LW6429 | *arIs37[secreted CC::gfp] I; cup-5(ar465) III; smoc-1(jj411[smoc-1(TY)::2xflag]) V; sma-9(cc604) X isolate #2* |

| | |
|---|---|
| LW6395 | *smoc-1(jj441[smoc-1(EC)::2xflag]) V* |
| LW6425 | *arIs37[secreted CC::gfp] I; smoc-1(jj441[smoc-1(EC)::2xflag]) V; sma-9(cc604) X isolate #1* |
| LW6427 | *arIs37[secreted CC::gfp] I; smoc-1(jj441[smoc-1(EC)::2xflag]) V; sma-9(cc604) X isolate #2* |
| LW6427 | *arIs37[secreted CC::gfp] I; cup-5(ar465) III; smoc-1(jj441[smoc-1(EC)::2xflag]) V; sma-9(cc604) X isolate #3* |
| LW6426 | *ccIs4438[intrinsic CC::gfp] III; smoc-1(jj441[smoc-1(EC)::2xflag]) V; sma-9(ok1628) X isolate #1* |
| LW6428 | *ccIs4438[intrinsic CC::gfp] III; smoc-1(jj441[smoc-1(EC)::2xflag]) V; sma-9(ok1628) X isolate #2* |
| **Strains overexpressing SMOC-1 truncations** | |
| LW6053 | *jjEx6053[pMSD44.4(smoc-1(TY)::2xflag) + LiuFD290(ttx-3p::RFP)]* |
| LW6054 | *jjEx6054[pMSD44.4(smoc-1(TY)::2xflag) + LiuFD290(ttx-3p::RFP)]* |
| LW6051 | *jjEx6051[pMSD44.4(smoc-1(TY)::2xflag) + LiuFD290(ttx-3p::RFP)]; smoc-1(tm7125) V* |
| LW6092 | *jjEx6092[pMSD44.4(smoc-1(TY)::2xflag) + LiuFD290(ttx-3p::RFP)]; smoc-1(tm7125) V* |
| LW6089 | *jjEx6089[pMSD44.4(smoc-1(TY)::2xflag) + LiuFD290(ttx-3p::RFP)]; arIs37[secreted CC::gfp] I; cup-5(ar465) III; smoc-1(tm7125) V; sma-9(cc604) X* |
| LW6090 | *jjEx6090[pMSD44.4smoc-1(TY)::2xflag) + LiuFD290(ttx-3p::RFP)]; arIs37[secreted CC::gfp] I; cup-5(ar465) III; smoc-1(tm7125) V; sma-9(cc604) X* |
| LW6057 | *jjEx6057[pMSD45.4(smoc-1(EC)::2xflag) + LiuFD290(ttx-3p::RFP)]* |
| LW6058 | *jjEx6058[pMSD45.4(smoc-1(EC)::2xflag) + LiuFD290(ttx-3p::RFP)]* |
| LW6052 | *jjEx6052[pMSD45.4(smoc-1(EC)::2xflag) + LiuFD290(ttx-3p::RFP)]; smoc-1(tm7125) V* |
| LW6093 | *jjEx6093[pMSD45.4(smoc-1(EC)::2xflag) + LiuFD290(ttx-3p::RFP)]; smoc-1(tm7125) V* |
| LW6117 | *jjEx6117[pMSD45.4(smoc-1(EC)::2xflag) + LiuFD290(ttx-3p::RFP)]; arIs37[secreted CC::gfp] I; cup-5(ar465) III; smoc-1(tm7125) V; sma-9(cc604) X* |
| LW6118 | *jjEx6118[pMSD45.4(smoc-1(EC)::2xflag) + LiuFD290(ttx-3p::RFP)]; arIs37[secreted CC::gfp] I; cup-5(ar465) III; smoc-1(tm7125) V; sma-9(cc604) X* |
| **Strains overexpressing SMOC-1 with various point mutations** | |
| LW6059 | *jjEx6059[(pMSD46.1(smoc-1(jj65 C210Y)::2xflag) + LiuFD290(ttx-3p::RFP)]* |
| LW6060 | *jjEx6060[(pMSD46.1(smoc-1(jj65 C210Y)::2xflag) + LiuFD290(ttx-3p::RFP)]* |
| LW6077 | *jjEx6077[pMSD47.7(smoc-1(jj85 E105K)::2xflag) + LiuFD290(ttx-3p::RFP)]* |
| *LW6083* | *jjEx6083[pMSD46.1(smoc-1(jj65 C210Y)::2xflag) + LiuFD290(ttx-3p::RFP)]; smoc-1(tm7125) V* |
| *LW6084* | *jjEx6084[pMSD46.1(smoc-1(jj65 C210Y)::2xflag) + LiuFD290(ttx-3p::RFP)]; smoc-1(tm7125) V* |
| *LW6085* | *jjEx6085[pMSD47.7(smoc-1(jj85 E105K)::2xflag) + LiuFD290(ttx-3p::RFP)]; smoc-1(tm7125) V* |
| *LW6086* | *jjEx6086[pMSD47.7(smoc-1(jj85 E105K)::2xflag) + LiuFD290(ttx-3p::RFP)]; smoc-1(tm7125) V* |
| **Strains carrying endogenously tagged HA::DBL-1** | |
| LW5863 | *dbl-1(jj307[HA::dbl-1 active domain]) V* |
| LW5864 | *dbl-1(jj308[HA::dbl-1 active domain]) V* |

| | |
|---|---|
| LW5865 | *dbl-1(jj309[HA::dbl-1 active domain]) V* |
| LW5933 | *dbl-1(jj308[HA::dbl-1]) V; ccIs4438[intrinsic CC::gfp] III; sma-9(cc604) X, isolate #1* |
| LW5934 | *dbl-1(jj308[HA::dbl-1]) V; ccIs4438[intrinsic CC::gfp] III; sma-9(cc604) X, isolate #2* |
| **Strains overexpressing tagged HA::DBL-1** | |
| LW6448 | *jjIs6448[pTYC3(dbl-1p::HA::dbl-1 gDNA::dbl-1 3'UTR) + LiuFD188 (myo-2p::mCherry)] x0* |
| LW6530 | *jjIs6448[pTYC3(dbl-1p::HA::dbl-1 gDNA::dbl-1 3'UTR) + LiuFD188 (myo-2p::mCherry)] x3* |
| **Strains overexpressing both SMOC-1::2xFLAG and HA::DBL-1** | |
| LW6592 | *jjIs5799[pMSD35.7(smoc-1p::smoc-1::2xflag) + LiuFD290(ttx-3p::RFP)]; jjIs6448[pTYC3(dbl-1p::HA::dbl-1 gDNA::dbl-1 3'UTR) + LiuFD188 (myo-2p::mCherry)], isolate #1* |
| LW6593 | *jjIs5799[pMSD35.7(smoc-1p::smoc-1::2xflag) + LiuFD290(ttx-3p::RFP)]; jjIs6448[pTYC3(dbl-1p::HA::dbl-1 gDNA::dbl-1 3'UTR) + LiuFD188 (myo-2p::mCherry)], isolate #2* |
| **Strains overexpressing SMOC-1::2xFLAG in the *lon-2(e678)* null background** | |
| LW6611 | *jjIs5799[pMSD35.7(smoc-1p::smoc-1::2xflag) + LiuFD290(ttx-3p::RFP)]; lon-2(e678), isolate #1* |
| LW6612 | *jjIs5799[pMSD35.7(smoc-1p::smoc-1::2xflag) + LiuFD290(ttx-3p::RFP)]; lon-2(e678), isolate #2* |
